# Cell-type-specific DNA methylation dynamics in the prenatal and postnatal human cortex

**DOI:** 10.1101/2025.02.21.639467

**Authors:** Alice Franklin, Jonathan P. Davies, Nicholas E Clifton, Georgina E T Blake, Rosemary Bamford, Emma M Walker, Barry Chioza, Martyn Frith, Joe Burrage, Nick Owens, Shyam Prabhakar, Emma Dempster, Eilis Hannon, Jonathan Mill

## Abstract

The human cortex undergoes extensive epigenetic remodelling during development, although the precise temporal and cell-type-specific dynamics of DNA methylation remain incompletely understood. In this study, we profiled genome-wide DNA methylation across human cortex tissue from donors aged 6 post-conception weeks (pcw) to 108 years of age. We observed widespread, developmentally regulated, changes in DNA methylation, with pronounced shifts occurring during early- and mid-gestation that were distinct from age-associated modifications in the postnatal cortex. Using fluorescence-activated nuclei sorting (FANS), we optimized a protocol for the isolation of SATB2-positive neuronal nuclei, enabling the identification of cell-type-specific DNA methylation trajectories in developing neuronal and non-neuronal populations. Developmentally dynamic DNA methylation sites were significantly enriched near genes implicated in autism and schizophrenia, supporting a role for epigenetic dysregulation in neurodevelopmental disorders. Our findings underscore the prenatal period as a critical window of epigenomic plasticity in the central nervous system with important implications for understanding the genetic basis of neurodevelopmental phenotypes.

## INTRODUCTION

Development of the human cortex is a complex and highly orchestrated process underpinned by the temporally coordinated regulation of gene expression ^1^. Starting with the rapid proliferation of neural progenitor cells in the ventricular zone, these transcriptional programs drive key developmental processes including neurogenesis and synaptogenesis and act to regulate neuronal growth, migration and connectivity in the developing cortex. Importantly, dysregulation to these pathways has been implicated in a range of neurodevelopmental disorders including autism and schizophrenia ^2,3^. The cortex continues to develop throughout fetal life and into the postnatal period when synaptic pruning and myelination further refine neural circuits during childhood and adolescence ^4^.

Epigenetic modifications are essential for the dynamic regulation of gene function during cellular differentiation in the developing central nervous system. The most studied epigenetic mechanism is DNA methylation, which involves the addition of a methyl group to the fifth carbon of cytosine. DNA methylation is primarily thought to inhibit local gene expression by disrupting transcription factor binding and attracting methyl-binding proteins that promote chromatin compaction and gene silencing ^5^. However, its effects on transcription can vary depending on the genomic and cellular context. For instance, DNA methylation within the gene body is often linked to increased gene expression ^6^ and has been associated with other genomic processes, such as alternative splicing and promoter usage ^7^. There is growing recognition about the role of epigenetic processes in mediating transcriptional plasticity in the developing central nervous system ^8^. DNA methylation plays a crucial role in neurodevelopment, as evidenced by the dynamic expression of the *de novo* DNA methyltransferases DNMT3A and DNMT3B in the developing brain ^9^. Mutations in the methyl-CpG binding protein 2 (MECP2) gene, which regulates neuronal gene expression by interacting with methylated DNA, lead to severe neurodevelopmental deficits in humans ^10^. Moreover, the regulation of DNA methylation is known to be involved in key neurobiological and cognitive functions throughout life, including neuronal plasticity ^8^, memory formation and retention ^11^, and circadian rhythms ^12^. Current analyses of DNA methylation in the developing cortex have been undertaken using tissue from a small number of donors spanning a narrow range of developmental ages ^13,14^ and little is known about the extent to which these changes continue during later stages of fetal development and during postnatal life. Importantly, given the role of DNA methylation in establishing cellular identity, existing studies have not systematically explored developmental patterns of DNA methylation in specific cell populations.

In this study we quantified DNA methylation across the genome in fetal cortex samples, reporting dramatic changes in DNA methylation across development, which are specific to the early- and mid-gestational period and distinct to age-associated changes observed in late gestation and the postnatal cortex. We developed a fluorescence-activated nuclei sorting (FANS) protocol to isolate SATB2-positive nuclei and use this approach to identify cell-type-specific trajectories of DNA methylation associated with the development of neuronal and non-neuronal cell-types. Finally, we show that neurodevelopmentally dynamic DNA methylation sites are enriched in the vicinity of genes implicated by genetic studies of autism and schizophrenia, supporting a role for neurodevelopmental processes in these conditions. This is, to our knowledge, the most extensive study of DNA methylation across development of the human cortex and confirms the prenatal period as a time of considerable epigenomic plasticity.

## RESULTS

### Dramatic changes in DNA methylation occur during development of the human cortex

Our first analyses focused on characterizing changes in DNA methylation during early- and mid-gestation fetal cortex development. Using cortex tissue from 91 fetal donors (age range = 6 to 23 post-conception weeks (pcw), male = 45 (age range = 6 to 20 pcw), female = 46 (age range = 8 to 23 pcw), **Supplementary Table 1**), DNA methylation was quantified across the genome using the Illumina EPIC microarray (see **Methods**). After stringent pre-processing, our final dataset included DNA methylation data for 807,806 sites (789,981 autosomal sites and 17,626 on the X chromosome). Using an epigenetic clock calibrated specifically for fetal brain ^15^, we found the expected strong correlation between predicted and actual developmental age (corr = 0.942, **Supplementary Figure 1**). We fitted a linear model controlling for sex and experimental batch (see **Methods**) and identified widespread changes in DNA methylation associated with developmental age, finding 50,913 (6.30% of total) differentially methylated positions associated with cortex development (dDMPs) (50,329 (98.9%) autosomal, 572 (1.12%) X chromosome) at an empirically-derived experiment-wide significance threshold (P < 9x10^-8^) ^16^ (**Supplementary Table 2**). Effect sizes for dDMPs overlapping with the subset of sites (n = 20,502) also profiled in an independent cohort of fetal brain samples using the Illumina 450K array ^13^ were highly correlated across studies (corr = 0.824, **Supplementary Figure 2**). Consistent with our previous results, we found a small but highly significant enrichment of dDMPs becoming hypomethylated with cortex development (n = 28,780 (56.5%), p = 1.61x10^-191^) with the mean effect size being significantly greater for hypomethylated dDMPs than hypermethylated dDMPs (change in DNA methylation (%) per week: hypermethylated dDMPs = 1.03, hypomethylated dDMPs = -1.53, p < 1x10^-320^, **Supplementary Figure 3**). Reflecting this, global levels of DNA methylation, calculated by taking the mean across all autosomal DNA methylation sites included in the final dataset, became slightly, but significantly, lower across fetal cortex development (change in DNA methylation (%) per week = -0.0194, p = 3.33x10^-10^) (**Supplementary Figure 4**).

The top-ranked dDMP (cg08125539), located within the gene encoding insulin-like growth factor 2 mRNA binding protein 1 (*IGF2BP1*), was characterised by a rapid increase in DNA methylation in the developing cortex (increase in DNA methylation (%) per pcw = 4.46, SE = 0.183, p = 1.16x10^-40^, **Figure 1A**). *IGF2BP1* plays a critical role in the development of the human cortex, being highly expressed during early embryogenesis but silenced later in gestation and postnatally ^17^. The top-ranked dDMP becoming hypomethylated across cortex development was cg11884704, located within the gene encoding solute carrier family 25 member 25 (*SLC25A25*) (change in DNA methylation (%) per week = -2.71, SE = 0.114, p = 8.63x10^-40^, **Figure 1B**). Many of the dDMPs associated with cortex development were spatially co-located, clustering into developmentally differentially methylated regions (dDMRs) spanning multiple DNA methylation sites. Using *dmrff* ^18^, we identified 1,356 dDMRs (corrected p < 0.05, number of dDMPs ≥ 3) associated with cortical development, with a mean size of 369 bp, annotated to 2,292 genes (**Supplementary Table 3**). Many of the top-ranked dDMRs are proximal to genes with established roles in development and function of the cortex. For example, the most significant dDMR, characterized by a dramatic increase in DNA methylation across cortex development (chromosome 11: 31,846,414 - 31,849,262, mean change in DNA methylation (%) per week across region = 6.50, p < 1x10^-320^) spans 20 sites overlapping a CpG island in *PAX6* (**Figure 1C**), a gene encoding a potent brain-expressed transcription factor known to play a critical role in neurogenesis and cortical development ^19^.

**Figure 1.**
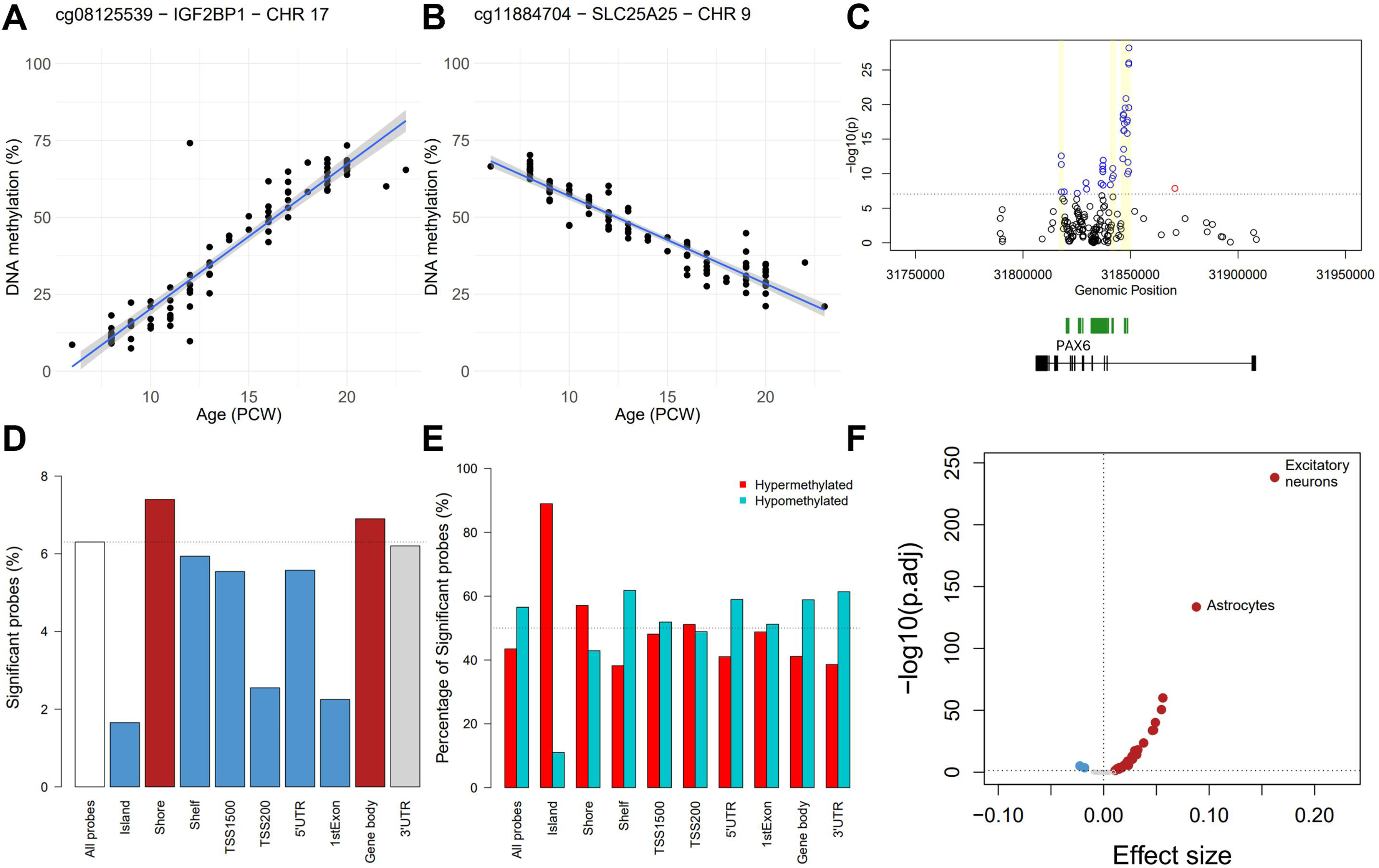
Development-associated changes in DNA methylation in the human fetal cortex. The top-ranked dDMPs demonstrating **A)** hypermethylation and **B)** hypomethylation over fetal cortex development. cg08125539, annotated to *IGF2BP1* on chromosome 17, is characterized by a significant increase in DNA methylation across prenatal cortex development (% increase in DNA methylation = 4.5% per week, p = 1.16x10^-40^). cg11884704, annotated to *SLC25A25* on chromosome 9, is characterized by a significant decrease in DNA methylation across prenatal cortex development (% decrease in DNA methylation = 2.7% per week, p = 8.63x10^-40^). Of note, the dramatic changes in DNA methylation at these sites are specific to the prenatal period (**Supplementary Figure 8**). **C)** dDMPs cluster into differentially methylated regions (DMRs) associated with cortex development (**Supplementary Table 3**). The *PAX6* gene on chromosome 11 contains 3 independent DMRs (yellow shaded regions), including the top-ranked DMR (mean change in DNA methylation (%) per week across region = 6.50, p < 1x10^-320^) spanning 20 sites overlapping an intragenic CpG island (green). Blue = dDMP with a positive effect size, red = dDMP with a negative effect size, black = non-significant site. Dotted line indicates genome-wide significance. **D)** Relative enrichment (red) and depletion (blue) of dDMPs across different CpG island and genic features. Compared to the frequency of dDMPs amongst all sites profiled in this study (white bar), there was a significant enrichment of dDMPs in CpG island shores and gene bodies but a significant depletion in other regions, most notably CpG islands (**Supplementary Table 5**). **E)** Proportion of hypermethylated (blue) and hypomethylated (red) dDMPs across CpG island and genic features. Despite the overall enrichment of hypomethylated dDMPs across all sites tested, there are substantial differences across genomic features with CpG islands and CpG island shores characterized by an enrichment of hypermethylated dDMPs (**Supplementary Table 6**). **F)** Enrichment of cortex dDMPs in cell-type-specific regions of open chromatin identified by scATAC-seq ^23^. Shown is a volcano plot of the relative effect size versus p-value from a logistic regression testing for an enrichment of dDMPs within cell-type-specific scATAC-seq peaks for 54 human fetal cell-types. Peaks for 31 fetal cell-types were significantly enriched for dDMPs, highlighting a general overlap with transcriptionally-active regions of the genome during development (red = significant enrichment, blue = significant depletion, gray = non-significant). The greatest enrichment was observed within ATAC-seq peaks specific to fetal excitatory neurons (odds ratio = 1.18, corrected p = 7.37x10^-239^) and astrocytes (odds ratio = 1.09, corrected p = 2.90x10^-134^).

### Developmental DMPs are enriched in specific genomic features and developmentally-active regions of open chromatin

Although dDMPs were distributed relatively equally across autosomal chromosomes, certain chromosomes were characterized by a relative enrichment or depletion (**Supplementary Figure 5** and **Supplementary Table 4**), most notably chromosome 19 (percentage of dDMPs = 3.49%, log odds ratio = -1.81, p = 2.16x10^-109^); of note, this chromosome has the highest gene density of any human chromosome ^20^. The distribution of dDMPs across genic features was more dramatically skewed (**Figure 1D** and **Supplementary Table 5**); for example there was a significant depletion of dDMPs in promoter regulatory regions including CpG islands (CGIs) (percentage of dDMPs = 1.65%, log odds ratio = -3.81, p < 1x10^-320^). In contrast, dDMPs were enriched in CGI shores (percentage of dDMPs = 7.40%, relative enrichment = 1.17, p = 1.74x10^-79^). dDMPs located in different genic features were also enriched for sites becoming either hypo- or hypermethylated during cortex development (**Figure 1E** and **Supplementary Table 6**). Despite the genome-wide enrichment of hypomethylated dDMPs, specific features were characterized by a dramatic enrichment of hypermethylated dDMPs including CGIs (hypermethylated DMPs = 89.0%, hypomethylated dDMPs = 11.0%, p < 1x10^-320^), and CGI shores (hypermethylated dDMPs = 57.1%, hypomethylated dDMPs = 42.9%, p = 1.50x10^-49^). These patterns resemble those seen in the development of other tissues ^21^ and reflect the observation that CpG islands typically exhibit low DNA methylation levels early in development, with tissue-specific DNA methylation patterns becoming established during the prenatal period ^22^. Finally, using publicly available single-cell ATAC-seq data generated from multiple human fetal tissues ^23^ we explored the extent to which autosomal dDMPs (n = 50,329) were enriched in regions of open chromatin associated with the development of different cell-types. dDMPs were significantly enriched in the top 10,000 ATAC-seq peaks identified in 31 out of 54 fetal cell-types tested, suggesting a broad overlap with transcriptionally-active regions of the genome during development (**Supplementary Table 7**). Strikingly however, the greatest enrichment was observed within ATAC-seq peaks specific to fetal excitatory neurons (odds ratio = 1.18, corrected p = 7.37x10^-239^) and astrocytes (odds ratio = 1.09, corrected p = 2.90x10^-134^) (**Figure 1F** and **Supplementary Figure 6**).

### Neurodevelopmental changes in DNA methylation are largely distinct to those occurring with age in the postnatal cortex

Given the major shifts in DNA methylation observed during cortex development, we were interested in characterizing the extent to which these changes were maintained across the life-course. We quantified DNA methylation in cortex tissue from postnatal donors (n = 673, age range = 0 to 104 years, **Supplementary Table 1**) in addition to a small number of late fetal cortex samples (n = 4, age range = 26 to 33 pcw, **Supplementary Table 1**). As expected, chronological age was strongly correlated with age estimates derived from DNA methylation data using both a pan-tissue epigenetic clock ^24^ (corr = 0.875, **Supplementary Figure 7A**) and a clock trained specifically on postnatal human cortex ^25^ (corr = 0.941, **Supplementary Figure 7B**). Overall, DNA methylation at dDMPs was dramatically more variable across early- and mid-fetal cortex samples than adult cortex samples (**Figure 2A**) (mean variance in DNA methylation (%) across the top 10,000 dDMPs profiled in both fetal and adult cortex = 120 and 41.7 respectively, p < 1x10^-320^). We next tested the extent to which development-associated DMPs were also characterized by age-associated changes in DNA methylation in the postnatal cortex, finding that for the 41,518 dDMPs also tested in our postnatal cortex samples (81.5% of all dDMPs) there was minimal correlation in age effect size between prenatal and postnatal cortex (corr = -0.00925) (**Figure 2B**). DNA methylation at only a very small proportion of these sites (n = 1,003 (2.42%)) was significantly (p < 1.20x10^-6^) associated with age in postnatal samples, with 275 (27.4%) of these sites being characterized by age-associated changes in the *opposite* direction to those observed in the developing cortex (**Figure 2C-F**). Of note, dDMPs that become hypermethylated across cortex development showed a higher proportion of consistent postnatal age effects (n = 664, (89.4%)) compared to sites becoming hypomethylated across cortex development (n = 124 (24.6%)). For the majority of cortex dDMPs, mean levels of DNA methylation in adult cortex (n = 661, age range = 25 - 104 years (average age = 81.8 years)) were highly correlated to the mean DNA methylation of the oldest fetal samples (n = 12 donors, aged 20 - 23 pcw), and dramatically different to that of the youngest fetal samples (21 donors, aged 6 - 9 pcw) (**Figure 2G**). Indeed, the top hyper- and hypomethylated dDMPs (cg08125539 (annotated to *IGF2BP1)* and cg11884704 (annotated to *SLC25A25*), respectively) show little variation in DNA methylation level after birth (**Supplementary Figure 8**), exemplifying the relative stability of DNA methylation postnatally following the dramatic developmental shifts observed in the prenatal cortex.

**Figure 2.**
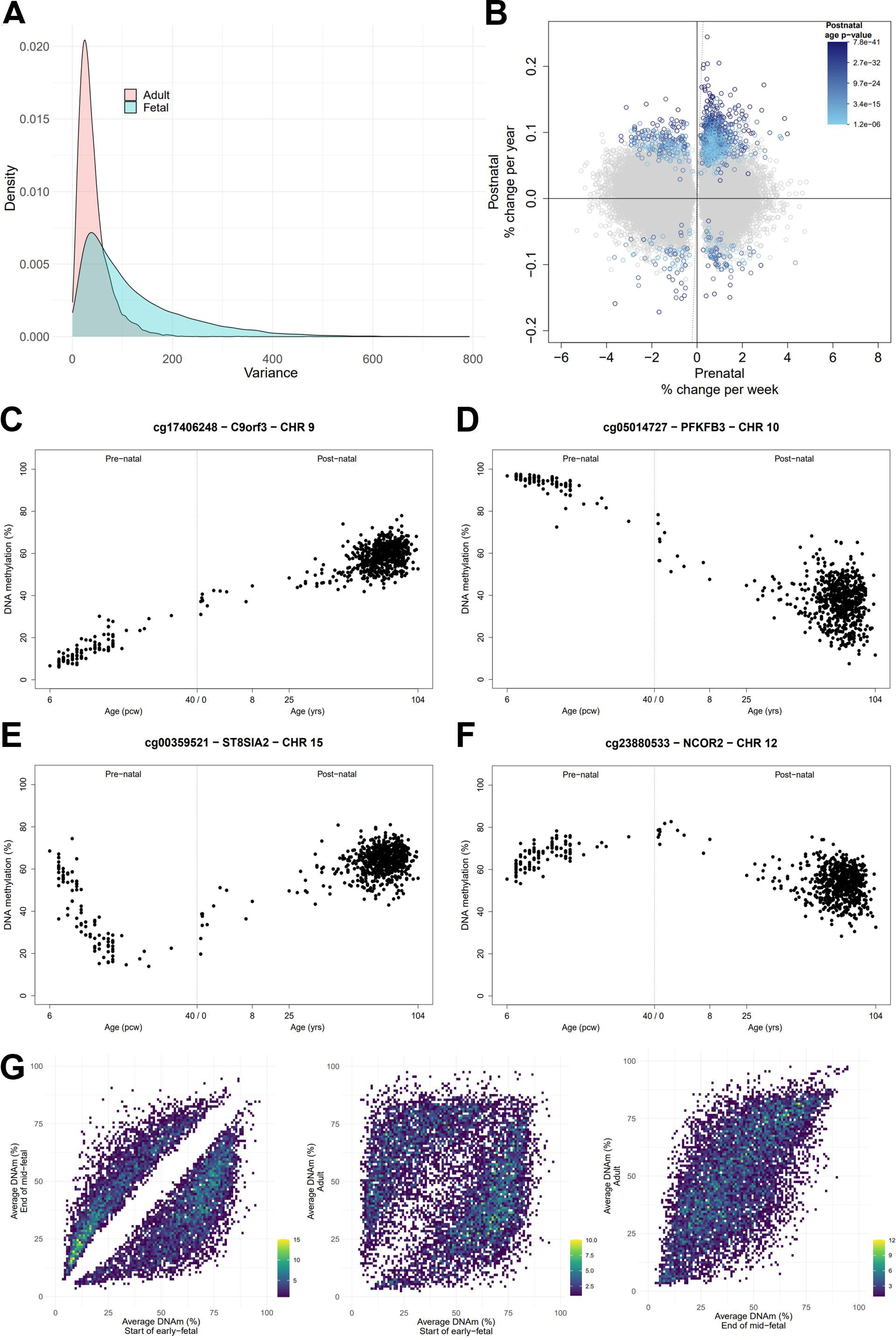
The majority of dDMPs are not characterized by age-associated changes in DNA methylation in the postnatal cortex. **A)** DNA methylation at dDMPs was dramatically more variable in fetal cortex samples than adult cortex samples. Shown is the distribution of variance in DNA methylation across the 10,000 most significant dDMPs in fetal samples (pink) (mean variance = 120, mean sd = 11% DNA methylation) compared to the variance in DNA methylation at the same sites in postnatal samples (blue) (mean variance = 41.7, mean sd = 6.46% DNA methylation). **B)** Comparison of age effect sizes between prenatal and postnatal cortex for the 41,518 dDMPs also measured in the postnatal samples. Each dot represents a cortex dDMP with significance for age in the postnatal cortex indicated by color (gray = not significantly associated with age in postnatal cortex). Examples of dDMPs showing **C)** a consistent age-associated increase in DNA methylation in both prenatal and postnatal cortex, **D)** a consistent age-associated decrease in DNA methylation in both fetal and postnatal cortex, **E)** a developmentally-associated decrease in DNA methylation in the prenatal cortex followed by an age-associated increase in DNA methylation in the postnatal cortex, and **F)** a developmentally-associated increase in DNA methylation in the prenatal cortex followed by an age-associated decrease in DNA methylation in the postnatal cortex. **G)** Comparisons of average (mean) DNA methylation at the 10,000 top-ranked dDMPs between the earliest fetal samples (n = 21, age range = 6 – 9 pcw), the eldest mid-fetal samples (n = 12, age range = 20 – 23 pcw) and adult samples (n = 661, age range = 25 - 104 years). Mean DNA methylation of the mid-fetal samples is most strongly correlated to postnatal samples (corr = 0.65) (right) than to early-fetal samples (corr = 0.36) (left).

### DNA methylation at a large proportion of sites changes non-linearly across cortex development

The regression model used to identify dDMPs was unable to distinguish between linear and nonlinear changes in DNA methylation in the developing cortex. Nonlinear changes could provide important insights into epigenetic switch-points during key stages of brain development ^26^ and therefore we sought to apply non-parametric modelling to identify nonlinear trajectories of DNA methylation. Using Gaussian process regression and model selection, we classified DNA methylation sites as constant, linear, or nonlinear throughout early- and mid-fetal cortex development. To reduce the chance of falsely identifying nonlinear patterns, we excluded non-variable probes, removed outlier samples, and focused on sites where DNA methylation changed within a biologically relevant timeframe (see **Methods**). Using this approach, we identified 73,035 sites (72,310 (99.0%) autosomal) that demonstrated high-confidence nonlinear changes in DNA methylation during the fetal period (**Supplementary Table 8**). This includes 23,252 (45.7%) of the dDMPs detected by our linear model (**Supplementary Figure 9A**) highlighting that the rate of change in DNA methylation at these sites is not consistent across development of the cortex (**Supplementary Figures 9B-C**). We applied weighted gene correlation network analysis (WGCNA) ^27^ to group autosomal sites characterized by shared nonlinear DNA methylation changes, finding six distinct modules (**Figure 3** and **Supplementary Table 8**). Gene ontology analysis of the genes annotated to sites in each module in turn revealed an enrichment of distinct pathways relevant to brain development and function (**Supplementary Table 9**). For example, with respect to all other sites on the array the turquoise module is enriched for genes involved in ‘regulation of cell differentiation’ (p = 6.13x10^-13^) and ‘mesenchyme development’ (p = 1.10x10^-7^), the ‘blue’ module for ‘neuron development’ (p = 1.68x10^-8^) and ‘gliogenesis’ function (p = 9.37x10⁻^7^), the ‘brown’ module for ‘neural tube closure’ (p = 1.88x10^-6^) and ‘neuron projection membrane’ (p = 4.49x10^-6^) and the ‘red’ module for ‘glial cell projection’ (p = 2.89x10^-8^) and ‘axonogenesis’ (p = 3.01x10^-6^).

**Figure 3.**
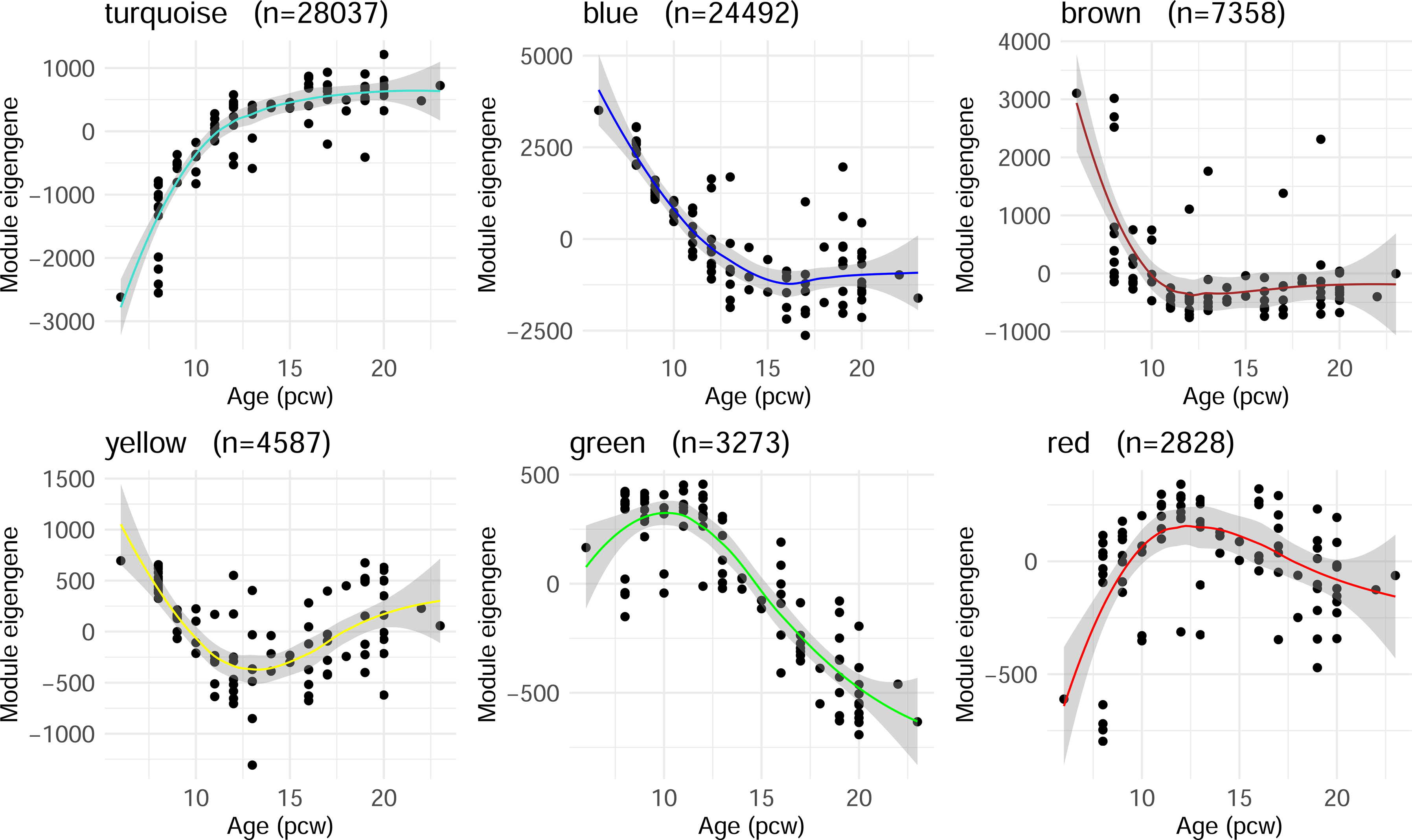
Nonlinear trajectories of DNA methylation in the developing cortex. Nonlinear DNA methylation sites identified through Gaussian process regression modelling were clustered into distinct modules using WGCNA (**Methods**). For each module, the eigengene (first principal component) is plotted against fetal age in post-conception weeks (pcw). The number of nonlinear sites assigned to each module is indicated in the subtitles, with hub DNA methylation sites for each module shown in **Supplementary Figure 25**.

### Developing a method to profile DNA methylation in excitatory neurons in the developing human cortex

Different neural cell-types are characterized by distinct transcriptional trajectories across development of the human cortex ^28^. As the analysis of bulk cortex tissue precludes the ability to identify changes in DNA methylation occurring in specific cell-types and might be confounded by the shifts in cell-type proportions occurring during this period ^29^, we next sought to isolate nuclei from developing neurons and non-neuronal cells prior to methylomic profiling. The nuclear marker NeuN (encoded by the *RBFOX3* gene) is widely used to label neuronal nuclei prior to fluorescent-activated nuclei sorting (FANS) and genomic profiling in postnatal human cortex ^30^ and we attempted to use this approach to profile DNA methylation in cortical NeuN+ and NeuN-nuclei populations obtained from donors spanning the life-course (see **Methods**). Despite NeuN robustly identifying discrete neuronal nuclei populations in late fetal and postnatal cortex samples, we found that it was not a reliable nuclear neuronal lineage marker in early- and mid-fetal donors. Although *RBFOX3* is expressed in postmitotic neurons and plays a role in neuron differentiation ^31^, interrogation of RNA-seq data generated by the BrainSeq Consortium ^32^ shows its expression is dramatically lower in fetal cortex compared to *SATB2* (Special AT-rich sequence-Binding protein 2), a potent transcription factor that drives early neurogenesis in the developing cortex and is postnatally expressed in excitatory neurons ^33^ (t = -14.562, p < 1x10^-320^, **Supplementary Figure 10**). We therefore optimized a method to immunolabel SATB2+ and SATB2-nuclei prior to FANS purification (see **Methods**), using snRNA-seq to confirm the coexpression of *RBFOX3* and *SATB2* in postnatal excitatory neurons (**Supplementary Figure 11**) and the higher expression of *SATB2* compared to *RBFOX3* in SATB2+ nuclei isolated from the fetal cortex (**Supplementary Figure 12**). Our protocol for the isolation of SATB2+ nuclei from fetal cortex is available as a resource to the community on Protocols.io (https://dx.doi.org/10.17504/protocols.io.n92ldz9d8v5b/v1).

### Differences in neuronal DNA methylation patterns between fetal and adult cortex

We used FANS to isolate neuron-enriched (SATB2+) and non-neuronal-enriched (SATB2-) nuclei populations from a subset (n = 37) of prenatal cortex tissue samples (**Supplementary Table 1**), profiling DNA methylation in each purified fraction as previously described (see **Methods**). These data were combined with existing DNA methylation data generated on neuron enriched (NeuN+), oligodendrocyte-enriched (SOX10+) and microglia-enriched (IRF8+ and NeuN-/SOX10-)) nuclei from 212 postnatal cortex samples by our group resulting in a cell-type-specific DNA methylation dataset from 259 donors aged 8 pcw to 108 years old (**Supplementary Table 1**). Principal component analysis (PCA) on the top 10,000 most variable autosomal sites calculated across the entire dataset highlighted that postnatal cell-type explained the majority of variance in DNA methylation (PC1, 71.5% of the variance), with developmental stage contributing significantly to both PC2 (17.1% of the variance) and PC3 (6.47% of the variance) (**Figure 4A**). We identified 6,531 DMPs between neuronal and non-neuronal nuclei isolated from early-/mid-fetal cortex (n = 37 donors, 8 - 20pcw). DNA methylation at these sites perfectly discriminated between neuronal and non-neuronal nuclei in the postnatal cortex (**Figure 4B**) and with much larger effect sizes, highlighting that cell-specific DNA methylation profiles are established early in development prior to terminal differentiation. Overall, we identified a much more striking difference between neuronal and non-neuronal nuclei in adult cortex (n = 212 donors, 18 - 108y, 453,675 DMPs) (**Supplementary Table 10**); although cell-type differences at these sites were significantly smaller in the fetal cortex (mean absolute effect size (%): fetal = 1.61, adult = 14.4, p < 1x10^-320^), it is striking that they still discriminate between neuronal and non-neuronal nuclei during early fetal development (**Figure 4C**). Despite an overall positive correlation of neuronal vs non-neuronal DNA methylation differences between fetal and adult cortex across all sites tested (corr = 0.230, **Supplementary Figure 13A**, **Supplementary Table 10**), many cell-type DMPs were characterized by developmental-stage-specific effects (**Supplementary Figures 13B-C**). Of particular interest were 1,427 sites that showed significantly different levels of DNA methylation between neuronal and non-neuronal nuclei in both fetal cortex and adult cortex, but with an opposite direction of effect (**Figure 4D**). These results suggest that whilst neuron-specific patterns of DNA methylation are established early in development, there are some notable shifts in patterns of cell-type-specific DNA methylation during development of the human cortex. Of note these differences are predominantly driven by changes in DNA methylation in neurons and not non-neuronal cells.

**Figure 4.**
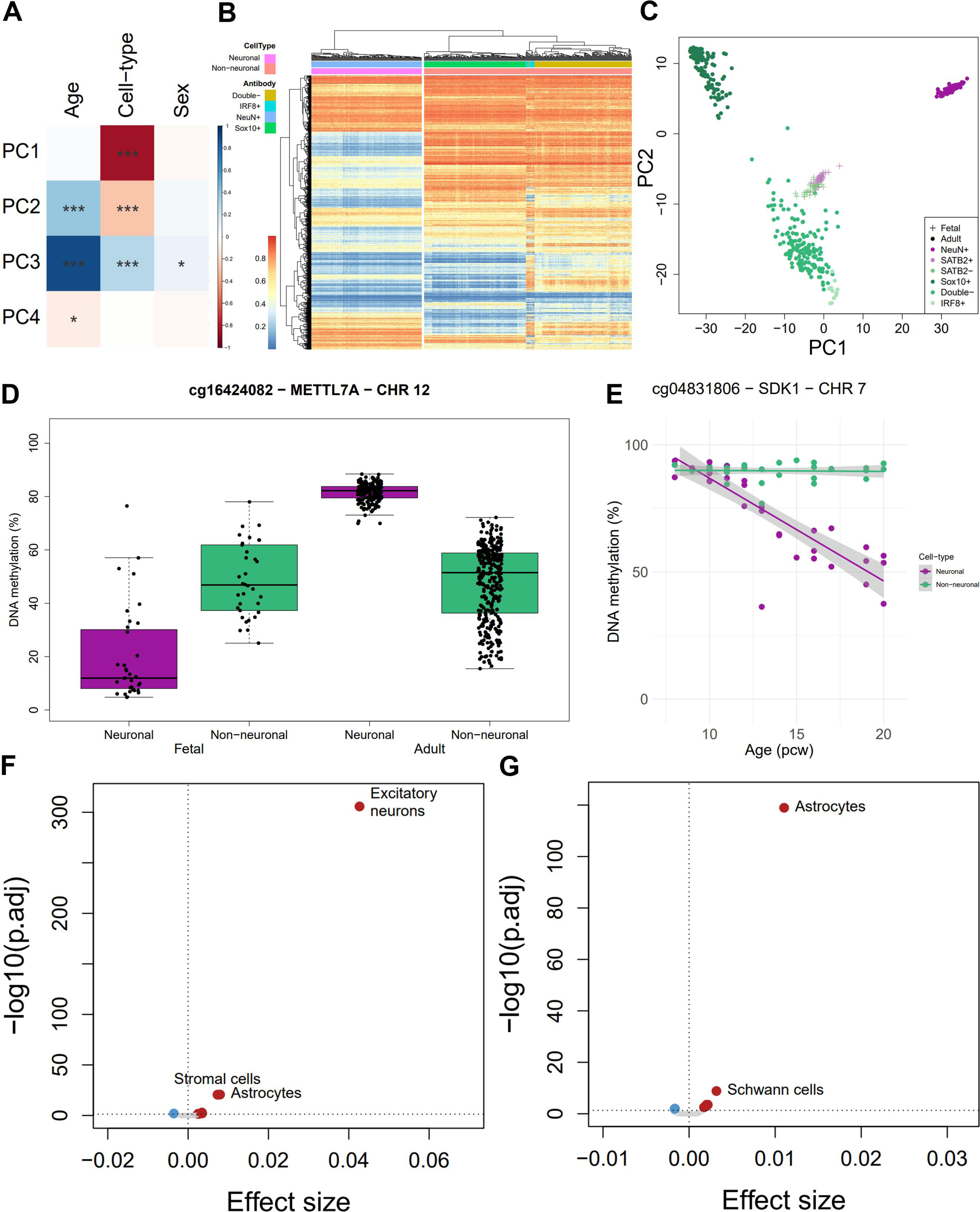
Cell-type-specific changes in DNA methylation across cortex development and postnatal life. **A)** Principal component analysis (PCA) on the top 10,000 most variable autosomal DNA methylation sites across FANS-isolated nuclei populations from prenatal and postnatal cortex samples (n = 259 donors). The top 4 principal components (PCs) correlated against age, cell-type and sex (p < 0.05 (*), p < 0.001 (***)). **B)** Heatmap of DNA methylation in cell-types isolated from adult cortex at the 6,531 DMPs between neuronal and non-neuronal nuclei identified in fetal cortex. **C)** PCA on the top 10,000 most variable autosomal sites in FANS-isolated nuclei populations from adult cortex (n = 212 donors). Using the adult PCA loadings, fetal samples were projected onto the same PCA space, showing a clear but attenuated difference between cell-types in fetal cortex. Colors indicate the antibody labels used to isolate cortical nuclei. Colors are grouped into neuronal (purple shades) and non-neuronal (green shades) cell-types. **D)** cg16424082, annotated to *METTL7A* on chromosome 12, is characterized by developmental-stage-specific differences in DNA methylation between neuronal and non-neuronal nuclei. **E)** DNA methylation at cg04831806 annotated to *SDK1* on chromosome 7, a gene with a key role in neurodevelopment ^34^, is characterized by a highly-significant interaction between cell-type and developmental age (neuronal % change in DNA methylation per week = -4.25, p = 6.76x10^-13^). **F)** Enrichment of neuronal dDMPs in cell-type-specific regions of open chromatin in the developing cortex identified by scATAC-seq ^23^. Neuron-specific autosomal dDMPs (n = 1,596) are most strongly enriched in peaks specific to excitatory neurons. **G)** Enrichment of non-neuronal dDMPs in cell-type-specific regions of open chromatin in the developing cortex identified by scATAC-seq ^23^. Non-neuronal-specific autosomal dDMPs (n = 548) are most strongly enriched in peaks specific to astrocytes and Schwann cells. Color indicates direction of effect: red = significant enrichment, blue = significant depletion, gray = non-significant.

### Distinct neuronal and non-neuronal trajectories of DNA methylation across cortex development

We used a linear model, adjusting for sex and experimental batch, to characterize changes in DNA methylation in neuronal and non-neuronal nuclei isolated from samples spanning early- to mid-fetal cortex development (n = 37 donors, **Supplementary Table 1**). Effect sizes for bulk cortex dDMPs that were also tested in FANS-purified nuclei (n = 42,114) showed a high correlation with those in both neuronal (correlation = 0.923) and non-neuronal (correlation = 0.870) samples (**Supplementary Figure 14**). Although the power to detect experiment-wide significant developmental changes was limited within each nuclei population given the relatively small sample size, we identified 1,872 neuronal and 820 non-neuronal dDMPs (p < 9x10^-8^) (**Supplementary Figure 15**, **Supplementary Table 11**). Of note, 4,088 of the dDMPs identified in the bulk cortex showed significant developmental changes in neurons (p < 1.19x10⁻⁶, corrected for 42,114 dDMPs also tested in FANS-isolated nuclei), with 1,820 dDMPs showing significant developmental changes in non-neuronal nuclei (**Supplementary Figure 16**, **Supplementary Figure 17**). Although there was a moderate correlation in developmental age effect sizes between the two cellular populations (corr = 0.564), many of the dDMPs identified in neuronal nuclei were clearly distinct to those identified in non-neuronal nuclei, in contrast to non-neuronal dDMPs which tended to be shared across both cell-types (correlation between cell-types: neuronal dDMPs = 0.634, non-neuronal dDMPs = 0.873, **Supplementary Figure 18**). There were some striking examples of cell-type-specific developmental effects; using an interaction model between age and cell-type we identified 744 sites at which the developmental change in DNA methylation was significantly different between neuronal and non-neuronal nuclei (p < 9x10^-8^) (**Supplementary Figure 19**, **Supplementary Table 11**). The most notable difference was at a site annotated to *SDK1* (cg04831806: interaction p = 2.97x10^-19^, neuronal effect = -4.25 (p = 6.76x10^-13^), non-neuronal effect = -0.0984 (p = 0.602)) encodes an immunoglobulin with a critical role in neuronal development and synapse formation ^34^ (**Figure 4E**), with other striking interaction effects seen for sites annotated to *SYT1*, *PEX14*, *OAT* and *GJA1* (**Supplementary Figure 19**). Gene ontology analysis of genes annotated these sites revealed a significant enrichment of functional pathways related to the specialisation of neuronal cell function, and especially pathways related to synapse development and signalling (**Supplementary Table 12**). We used publicly available single-cell ATAC-seq data from multiple human fetal tissues ^23^ to test whether neuron-specific (n = 1,596) and non-neuron-specific (n = 548) autosomal dDMPs were enriched in regions of open chromatin associated with the development of specific cell-types. Neuron-specific dDMPs were significantly and specifically enriched in excitatory neuron ATAC-seq peaks (odds ratio = 1.04, Bonferroni-adjusted p = 1.73x10^-306^) with no such enrichment observed for non-neuron-specific dDMPs (odds ratio = 0.999, Bonferroni-adjusted p = 1). In contrast, non-neuronal-specific dDMPs were enriched in peaks defining cell-types including astrocytes (odds ratio = 1.01, Bonferroni-adjusted p = 1.15x10^-119^) and Schwann cells (odds ratio = 1.003, Bonferroni-adjusted p = 1.47x10^-9^) (**Figure 4F-G**, **Supplementary Figure 20**, **Supplementary Table 13**, and **Supplementary Table 14**).

### Genes implicated in autism and schizophrenia are enriched for neurodevelopmental changes in DNA methylation

Autism and schizophrenia are highly heritable neurodevelopmental conditions hypothesized to arise from differences in early cortex development ^35^, and recent exome- and whole genome-sequencing studies have identified a number of rare but highly-penetrant mutations associated with both conditions. We explored the extent to which DNA methylation sites annotated to both high confidence autism genes (n = 233 ‘Category 1’ autism genes from the (Simons Foundation Autism Research Initiative (SFARI) Gene database ^36^) and schizophrenia genes identified from exome-sequencing (n = 32 genes identified by the SCHEMA consortium ^37^) (**Supplementary Table 15**) were enriched for dDMPs (see **Methods**). Amongst autism and schizophrenia genes there was a significant enrichment of loci associated with at least one bulk cortex dDMP compared to the background rate of 43.7% for all genes annotated to DNA methylation sites (SFARI autism and SCHEMA schizophrenia genes combined = 69.8% (p = 2.96x10^-3^), SFARI autism genes = 70.4% (p = 8.38x10^-4^), SCHEMA schizophrenia genes = 64.5% (p = 0.270), **Supplementary Figure 21, Supplementary Table 16**). Of note, compared to the background rate of genes associated with dDMPs in neuronal (3.95%) and non-neuronal (1.89%) nuclei, SCHEMA schizophrenia genes were significantly enriched for genes annotated to dDMPs in both cell-types (neuronal: 12.9%, p = 1.30x10^-5^; non-neuronal: 9.68%, p = 2.98x10^-4^). In contrast, SFARI autism genes were enriched specifically for developmental changes in neuronal nuclei (15.7%, p = 4.35x10^-6^) but not non-neuronal nuclei (6.09%, p = 0.0962) (**Supplementary Figure 22, Supplementary Table 17**, **Supplementary Table 18**).

Both autism and schizophrenia also have a large polygenic component, and we next examined the enrichment of dDMPs in genomic regions containing common disease variants identified by genome-wide association studies (GWAS) using MAGMA gene set analysis ^38^ with the most recent GWAS results for autism ^39^ and schizophrenia ^40^ (see **Methods**). For schizophrenia we observed an overall enrichment amongst bulk cortex dDMPs (effect size = 0.0496, p = 7.43x10^-3^). Cell-type-specific analyses suggest this association is driven by neuronal dDMPs (effect size = 0.0873, SE = 0.0446, p = 0.0250) and not by non-neuronal dDMPs (effect size = -0.0944, SE = 0.0613, p = 0.938) (**Supplementary Figure 23**). No significant MAGMA enrichment was observed for autism (**Supplementary Figure 23**), likely reflecting the limited power of the current autism GWAS.

## DISCUSSION

In this study we characterised changes in cortical DNA methylation during early- and mid-fetal development, exploring the extent to which these trajectories continued in the postnatal cortex and identifying differences in developmental patterns of DNA methylation between neuronal and non-neuronal cell-types. We identified dramatic changes in DNA methylation associated with cortex development, with the majority of these effects specific to the prenatal period and distinct to age-associated changes in the postnatal cortex. We developed a novel fluorescence-activated nuclei sorting protocol to isolate SATB2+ neurons from the developing cortex and identify trajectories of DNA methylation associated with the development of neuronal and non-neuronal cell-types. Finally, we demonstrated an enrichment of developmentally dynamic DNA methylation sites annotated to genes implicated in autism and schizophrenia, supporting a role for neurodevelopmental processes in these conditions. This is, to our knowledge, the most extensive study of DNA methylation across the development of the human cortex and confirms the prenatal period as a time of considerable epigenomic plasticity in the human brain.

Several key findings emerge from our analyses. First, we show that the distribution of developmentally-associated differentially methylated positions differs across genomic regions, being depleted in CpG islands (likely due to their association with stably-expressed housekeeping genes ^41^) and enriched in regions of open chromatin identified in scATAC-seq analyses of fetal tissue development. Furthermore, although there was an overall enrichment of dDMPs becoming hypomethylated during development, in some genomic features the converse was true; for example, ∼90% of the dDMPs annotated to CpG islands became hypermethylated over prenatal development. This supports observations from previous studies of changes in DNA methylation during early- and mid-gestation in brain and other tissues ^13,21,22,42^ and reflects the role of CpG island methylation in mediating tissue-specific transcriptional programs during development ^43^.

Second, the dramatic changes in DNA methylation observed in the developing cortex were largely specific to the prenatal period. These changes were distinct from the age-related changes seen in the postnatal cortex, and DNA methylation at dDMPs remained relatively stable after birth. Among the small subset (∼2%) of dDMPs that did show significant postnatal changes in DNA methylation, many exhibited changes in the opposite direction to those seen across development of the fetal cortex. Specifically, sites that were hypomethylated during the prenatal period often became significantly hypermethylated postnatally, which is consistent with previous findings^42^. These results confirm that the prenatal period is a time of intense epigenetic activity in the developing cortex, with more extensive and larger DNA methylation changes compared to the postnatal period.

Third, our findings demonstrate that DNA methylation changes during cortex development are not uniformly linear. Instead, we identified distinct clusters of DNA methylation sites that exhibit nonlinear changes. These nonlinear changes could provide important insights into epigenetic milestones during key stages of brain development, many of which could be missed by standard linear regression approaches. Using a non-parametric Gaussian process regression model, we identified thousands of sites whose DNA methylation changed nonlinearly during fetal cortex development. Clustering these sites into six distinct co-methylation modules revealed shared nonlinear trajectories and highlighted clear switch-points in DNA methylation during development, particularly between 12 and 15 post-conception weeks. For example, nonlinear sites in the “blue” module were characterised by a rapid initial decrease in DNA methylation, plateauing around 15 pcw; pathway analysis of the genes annotated to these sites revealed a significant enrichment of genes involved in gliogenesis, which is understood to emerge around 15 pcw in humans ^44^. Additionally, sites in the “brown” module exhibited a rapid decrease in DNA methylation in the youngest samples and were annotated to genes enriched in pathways related to neural tube closure which occurs around the fourth post-conception week ^44^. Collectively, these findings highlight the utility of using nonlinear models to identify distinct patterns of DNA methylation in the developing fetal cortex that may be indicative of developmental switch-points that would not have been detected using a standard linear model.

Fourth, we find that there are distinct patterns of DNA methylation in developing neuronal and non-neuronal cell-types. The period of cortical development is marked by rapid neurogenesis and the terminal differentiation of excitatory neurons which arise from radial glial progenitor cells in the ventricular and subventricular zones, with this process starting ∼7 pcw and peaking between 12 and 20 pcw ^44,45^. To explore DNA methylation changes accompanying neurogenesis we optimized a method to separate neuronal and non-neuronal nuclei, identifying SATB2 as a robust marker of prenatal neurons. Our analysis revealed that cell-type differences accounted for a large proportion of the variation in DNA methylation, with consistent differences between neuronal and non-neuronal cells in both fetal and adult samples. Our analyses highlighted cell-type-specific DNA methylation trajectories during cortex development, with changes in bulk cortex primarily reflecting those occurring in developing neurons. Of note, analysis of a fetal scATAC-seq dataset revealed a striking enrichment of neuron-specific dDMPs in regions of open chromatin associated with the development of excitatory neurons and an enrichment of non-neuronal-specific dDMPs in regions of open chromatin associated with glial cell types (astrocytes and Schwann cells). These results confirm the interplay between DNA methylation and chromatin organisation during development ^46^ and suggest that dynamic changes in DNA methylation play an important role in establishing the broader cell-type-specific epigenetic landscape of the developing human cortex.

Finally, we also found evidence that dDMPs are enriched in sites annotated to genes associated with autism and schizophrenia, two highly heritable neurodevelopmental conditions that have been hypothesized to arise from differences in early cortex development ^35,47,48^. Genes robustly associated with both schizophrenia (from the SCHEMA consortium ^37^) and autism (from the SFARI gene database ^36^) were enriched for dDMPs, providing further evidence to support the notion that altered gene regulation during fetal brain development is involved in the etiology of these conditions. dDMPs were also enriched in the vicinity of common variants associated with schizophrenia by GWAS, although no enrichment was observed for common variants associated with autism, possibly reflecting the low power of the current autism GWAS. Our data support the hypothesis that a significant proportion of the genes harboring genetic variants that confer risk for neurodevelopmental conditions have regulatory effects that manifest during the development of the human cortex.

This study has several limitations that should be taken into consideration. First, legal restrictions on later-term abortions limited our access to cortical tissue from more advanced stages of fetal development. However, the relative stability of DNA methylation at dDMPs between the oldest fetal samples and adult cortex samples (**Figure 2A-B**) suggests that the magnitude of epigenetic changes during late pregnancy and early postnatal life is much smaller than those observed during early fetal development. Second, while the Illumina EPIC array enables precise quantification of DNA methylation at single-base resolution for many genes and CpG islands, it covers only a small proportion of CpG sites in the human genome, and these are not evenly distributed across all genomic features. As sequencing costs decline, future research should utilize sequencing-based technologies to comprehensively profile the epigenome across cortical development with larger sample sizes. Third, despite this being the largest study of cell-type-specific DNA methylation changes in human cortex development to date, the labor-intensive nature of isolating purified nuclei via FANS limited us to collecting neuronal and non-neuronal fractions from only a subset of donors, which somewhat reduced our power to detect dDMPs in these cell-types. Nevertheless, we observed strong concordance with bulk cortex data and clear evidence of cell-type-specific developmental trajectories. Fourth, our study could not distinguish between DNA methylation and its oxidized form, DNA hydroxymethylation, due to limitations of sodium bisulfite-based approaches ^49^. This distinction is critical, as DNA hydroxymethylation is abundant in the central nervous system and known to impact gene expression ^14^. Future studies should aim to differentiate these modifications using more refined techniques. Lastly, we did not obtain gene expression data from these samples, preventing direct conclusions about the transcriptional impact of the observed DNA methylation changes. However, by integrating our dDMPs with scATAC-seq peaks from fetal brain ^23^, we were able to show that the DNA methylation changes observed reflect broader shifts in gene regulation throughout cortical development.

In conclusion, our study uncovers highly dynamic and widespread changes in DNA methylation across the human cortex during development. These dramatic temporal shifts in DNA methylation are predominantly confined to the prenatal period, differing significantly from the more gradual, age-related changes observed in the postnatal cortex. Importantly, the enrichment of dDMPs near genes associated with schizophrenia and autism offers novel insights into the developmental mechanisms underlying these conditions. These findings provide insight into how early epigenetic modifications may mediate the onset of neurodevelopmental phenotypes, paving the way for future mechanistic research.

## MATERIALS & METHODS

### Human cortex tissue samples

Cortex tissue from prenatal and childhood donors was acquired from the Human Developmental Biology Resource (HDBR) (http://www.hdbr.org) and the MRC UK Brain Banks network (https://ukhealthdata.org/members/mrc-uk-brain-banks-network). Ethical approval for the HDBR was granted by the Royal Free Hospital research ethics committee under reference 08/H0712/34 and Human Tissue Authority (HTA) material storage license 12220; ethical approval for the MRC Brain Bank was granted under reference 08/MRE09/38. The age of donors was determined by Carnegie staging in the case of embryonic samples and foot and knee to heel length measurements for fetal samples. Apart from sex, no additional phenotypic or demographic information was available. A full overview of the samples used in this study is provided in **Supplemental Table 1**.

### DNA methylation profiling

Genomic DNA was isolated from each tissue sample using a standard phenol-chloroform extraction protocol and assessed for quality and purity using spectrophotometry. DNA methylation was quantified using the Illumina Infinium HumanMethylation EPIC array (Illumina Inc), which interrogates >850,000 DNA methylation sites across the genome. Briefly, the EZ-96 DNA Methylation-Gold kit (Zymo Research) was used for sodium bisulfite conversion prior to the quantification of DNA methylation using the Illumina EPIC array. All subsequent statistical analyses were performed in R version 3.6.0 unless otherwise stated. Raw Illumina EPIC data was processed using the *wateRmelon* R package as previously described ^50^. Briefly, DNA methylation data were loaded from raw IDAT files and processed through a standard quality control pipeline that includes the following steps: 1) Checking methylated and unmethylated intensities and excluding samples where this was < 500; 2) Calculating bisulfite conversion efficiency and excluding samples with median < 80%; 3) Principal component analysis of the X and Y chromosomes to confirm reported sex; 4) Confirming sample (un)relatedness using the 59 ‘rs’ SNP probes on the EPIC array; samples with correlation > 80% to another unrelated sample were excluded; 5) Detection of outlier samples with the *outlyx* function; 6) Using the *pfilter* function to identify and exclude samples with > 1% of probes with detection p > 0.05 and probes where > 1% of samples had a detection p > 0.05. 7) Exclusion of probes previously identified as cross-hybridising or polymorphic ^51^. 8) Normalisation with *dasen*. Data generated on the purified nuclei populations were confirmed through Principal Component Analysis (PCA) to cluster within cell-type and were normalised within each cell-type separately. EPIC array probes were annotated to genes according to the Illumina Infinium MethylationEPIC v1.0 B4 Manifest File (available at: https://emea.support.illumina.com/downloads/infinium-methylationepic-v1-0-product-files.html).

### Fluorescence-activated nuclei sorting (FANS) isolation of neuronal nuclei from fetal cortex

Neuronal and non-neuronal nuclei fractions were isolated from 38 fetal cortex tissue samples using a fluorescence-activated nuclei sorting (FANS) protocol optimised by our group. Briefly, following tissue homogenization and nuclei purification using sucrose gradient centrifugation we used a FACS Aria III cell sorter (BD Biosciences) to simultaneously collect populations of SATB2+ (AbCam, Cat No: ab196316, dilution: 1:1000) and SATB2-immunolabeled populations from fetal cortex prior to DNA methylation profiling. For each nuclei population, ∼200,000 nuclei were collected for extraction of genomic DNA using a standard phenol:chloroform extraction protocol and DNA methylation was profiled using the Illumina EPIC array as described above. Our detailed protocol detailing our SATB2+ FANS protocol is available on protocols.io ^52^ as a resource to the community.

### Generation of whole cortex life-course DNA methylation dataset

After quality control and normalisation, the fetal cortex DNA methylation dataset (n = 91, age range = 6 - 23 pcw) included 807,806 DNA methylation sites (789,981 autosomal). To generate a life-course dataset we incorporated DNA methylation data from 16 late-gestation fetal and child donors (age range = 26 pcw - 8 years) and 661 adult donors (age range = 25 - 104 years) using data previously reported by our group (Gene Expression Omnibus (GEO) accession number GSE197305 ^30^). The final bulk cortex life-course dataset included DNA methylation data from 768 donors (**Supplementary Table 1**) and included data for 41,518 of the 50,913 dDMPs identified in the fetal cortex.

### Generation of nuclei-sorted life-course DNA methylation dataset

We combined normalised neuronal (SATB2+) and non-neuronal (SATB2-) DNA methylation data derived from 37 early-/mid-fetal donors (neuronal n = 34, non-neuronal n = 33, age range = 8 - 20 pcw), 1 late-fetal donor (neuronal n = 1, non-neuronal n = 1, age = 28 pcw) and 9 child donors (neuronal n = 9, non-neuronal n = 9, age range = 0 - 8 years), generated using the FANS protocol described above, with data from 212 adult donors (neuronal (NeuN+) n = 178, non-neuronal (SOX10+, IRF8+ and SOX10-/NeuN-) n = 335, age range = 18 - 108 years) generated previously by our group as part of a large-scale study quantifying DNA methylation in human purified cortical nuclei ^53^. The final life-course dataset contained DNA methylation data for FANS-isolated nuclei populations from 259 donors (**Supplementary Table 1**) across 693,964 shared sites (679,917 autosomal).

### Identification of differentially methylated positions and regions associated with cortex development

We used a multiple linear regression model to test the association between developmental age in post-conception weeks (pcw) and DNA methylation, whilst controlling for sex and experimental batch. A site was considered to be a significant dDMP if its p-value surpassed an experiment-wide significance threshold of p < 9x10^-8^, previously determined to adequately control for the false positive rate of DNA methylation studies on the EPIC array ^16^. Regional analysis of dDMPs was performed using *dmrff* (https://github.com/perishky/dmrff) to identify differentially methylated regions (dDMRs). dDMRs were defined as regions with three or more dDMPs within a maximum window of 500 bp and reaching a Bonferroni-adjusted p < 0.05.

### Identification of cell-type specific trajectories across the life-course

Mixed-effects models were applied to test cell-type (neuronal vs non-neuronal) associations across the 579,541 DNA methylation sites shared between fetal and adult nuclei-sorted samples. Fixed effects included age (prenatal age in post-conception weeks; postnatal age in years), sex and array plate number and individual ID was included as a random effect to account for multiple measurements from each individual. SOX10+ and IRF8+ dummy variables were included as fixed effect covariates within the adult model to account for differences between the cell-type sub-populations within non-neuronal nuclei (SOX10+, IRF8+ and SOX10-/NeuN-). An ANOVA was used to determine probe significance by whether the addition of the variable of interest (cell-type) significantly improved the model (p < 9x10^-8^). Downstream analyses used the mixed-effects model coefficients.

### Genomic feature and chromosome enrichment

DNA methylation sites were annotated to genomic and CpG features according to the Illumina EPIC manifest. Enrichment of DMPs within features and chromosomes was tested with a chi-squared test. To determine if there was an overall enrichment of DMPs for a specific direction of effect (hypomethylated or hypermethylated), a one-tailed binomial test was used with probability of success equal to the proportion across all sites.

### Chromatin peak enrichment

We utilised publicly available scATAC-seq data generated by Domcke *et al.*^23^ to interrogate the top 10,000 most specific chromatin peaks for 54 fetal human cell-types (available at GEO accession number GSE149683, **Supplementary Table 19**). For each cell-type, autosomal DNA methylation sites were annotated with a binary variable indicating whether they resided within the genomic coordinates of each peak. We used logistic regression to test the association between a DNA methylation site being a DMP and being in a peak, whilst controlling for all other peaks for each cell-type.

### Enrichment of biological pathways and rare-variant gene sets for autism and schizophrenia

To test whether dDMPs were significantly associated with a gene set (biological pathway or gene list), we first annotated probes to genes as previously described. At the gene-level, we used logistic regression to test the relationship between DMP status (a binary variable of whether a gene contains at least one DMP) and gene set membership (whether the gene belongs to the gene set), whilst controlling for gene size (number of DNA methylation sites annotated to the gene). For Gene Ontology (GO) pathway analyses, the gene set consisted of the genes annotated to sites on the EPIC array that overlap with gene ontology pathways from the *GO.db* R package (v3.18). For enrichment of autism- and schizophrenia-related genes, the gene set consisted of published gene lists for each condition ^36,37^.

### Gene set enrichment of common variants associated with autism and schizophrenia

Enrichment analyses with respect to common genetic variants associated with autism ^39^ and schizophrenia ^40^ were performed using MAGMA v1.10 ^38^. Non-variable DNA methylation sites and sites not annotated to a gene were excluded prior to analysis. GWAS-associated SNPs were subset to common variants (minor allele frequency (MAF) ≥ 1% and imputation information score > 0.8) and SNP-associated GWAS p-values were combined into gene-level p-values using a window of 35 kb upstream and 10 kb downstream of the gene, as recommended ^54^. One-tailed MAGMA gene set analysis tested the association between gene-level SNP p-values and DMP status, whilst controlling for gene size (number of DNA methylation sites annotated to the gene).

### Characterization of nonlinear trajectories during prenatal cortex development

To determine sites with nonlinear trajectories, we first applied additional filtering. A leave-one-out Z-score approach was used to detect and remove the contribution of outlier samples (those with Z-scores > 5 standard deviations from the mean). Non-variable DNA methylation sites (sites whose range of the middle 80% of values is < 5% DNA methylation), and constant sites (sites with DNA methylation consistently > 90% or < 10% for all samples), were also removed. Using https://github.com/owensnick/GPMethylation.jl in Julia v1.6.1, we performed exact Gaussian process regression and model selection to classify DNA methylation sites as either constant, linear or nonlinear (Matern 5/2 kernel). Hyperparameters were selected by optimization of the marginal likelihood. Classification was determined by considering the kernel that provided the greatest optimised marginal log-likelihood. Nonlinear sites (n = 175,419) were further refined by considering the log-likelihood ratio (LLR) of the nonlinear kernel against both the constant and linear kernels (**Supplementary Figure 24**). The timescale of a DNA methylation site (traditionally called length-scale) relates to the periodicity of the oscillation, measured in post-conception weeks (pcw). To determine the period of one complete oscillation, the timescale can be multiplied by 2π sqrt(⅗) (≈ 5), i.e. a timescale of 1 equates to a complete oscillation every ≈ 5 pcw. To focus on sites with strong evidence for nonlinear behavior with biologically meaningful timescales, sites with LLR < 2 and timescale < 10 were excluded, leaving 73,638 high-confidence nonlinear sites with timescales between 10.0 and 106 (**Supplementary Figure 24**). We used the weighted gene correlation network analysis (WGCNA) R package ^27^ to identify modules of co-methylated nonlinear sites. Using the *blockwiseModules* function, a signed network was constructed using a soft threshold of 12, as recommended for this sample size ^55^. Representative module plots (**Figure 3**) were constructed by calculating the first principal component (eigengene) of each module. “Hub sites” were defined as sites whose DNA methylation pattern most highly correlated with the module eigengene (**Supplementary Figure 25**). Pathway analysis was performed on genes annotated to the nonlinear sites within each module separately, as described above, with the background gene list consisting of genes annotated to all other array sites.

### Single nuclei RNA-Seq (snRNA-Seq)

50mg of cortex tissue was homogenised using the recommended protocol from 10X Genomics (CG000124, Rev F) with a few minor amendments. Briefly, ribolock RNase-inhibitor (Thermo Scientific, EO0382) was added to the lysis buffer (0.4 U/ul) and staining buffer (0.2U/ul). Following antibody staining, nuclei were collected by FANS as described above. Nuclei were then spun and resuspended in staining buffer. Nuclei suspensions were assessed for the presence of debris and manually counted on a haemocytometer before proceeding with single-nucleus capture using the 10x Genomics Single-Cell 3′ technology. Targeted recovery of 3000 nuclei per sample was used. The 10x Chromium Next GEM Single Cell 3’ protocol (v3.1) was followed for expression library preparation from single nuclei. cDNA and final library quantification, quality control and fragment size determination was performed using the D5000 high sensitivity ScreenTape assay and reagents (Agilent technologies). Sequencing libraries were pooled and sequenced on an Illumina NovaSeq6000 sequencer. Library demultiplexing, generation of FASTQ files, alignment of reads and quantification of unique molecular identifiers (UMIs) was performed using CellRanger software with default parameters (10x Genomics, v3.1.0). A pre-mRNA reference file (GRCh38) was used to ensure unspliced intronic reads were captured. snRNA-seq data was analysed using the Seurat package in R (v4.1 and 3.6). Doublet identification was performed using the DoubletFinder R package. Cells were filtered prior to clustering on the basis of genes per cell (nFeature_RNA, data set specific) and percentage mitochondrial reads per cell (greater than 5% mitochondrial reads excluded). Data was log normalised and highly variable genes were selected. The data was then scaled (linear transformation) and variation owing to mitochondrial gene expression was regressed out. PCA was performed followed by clustering using a KNN graph. Dimensionality reduction was performed with t-distributed Stochastic Neighbour Embedding (tSNE) or uniform Manifold Approximation and Projection (uMAP). Clusters were manually annotated using analysis of differentially expressed marker genes from previous scRNA-seq studies ^56–58^ and plots were generated using the scCustomize package^59^.

## DATA AVAILABILITY

DNA methylation data generated as part of this study are publicly available via Gene Expression Omnibus (GEO) accession numbers GSE289191 (bulk fetal and child cortex) and GSE289184 (sorted fetal and child cortex). Bulk adult cortex data is available at GSE197305 ^30^ and sorted adult data is available at GSE279509 ^53^. Code relating to the analyses reported here can be found on GitHub (https://github.com/alicemfr/DevCortexDNAm/).

## ACKNOWLEDGEMENTS

This work was supported by grants from the Simons Foundation for Autism Research (SFARI) (grant number 573312, awarded to J.M. and grant number 809383, awarded to the APEX consortium), grants from the UK Medical Research Council (grant MR/R005176/1, awarded to J.M., and grant MR/W017156/1, awarded to N.C.) and the European Union’s Horizon Europe program (YOUTH-GEMs, grant agreement No. 101057182). The human prenatal material was provided by the Human Developmental Biology Resource (funded by MRC/Wellcome Trust grant (099175/ Z/12/Z) (https://www.hdbr.org). Sequencing infrastructure was supported by a Wellcome Trust Multi User Equipment Award (WT101650MA, awarded to J.M.) and Medical Research Council (MRC) Clinical Infrastructure Funding (MR/M008924/1, awarded to J.M.). This study was also supported by the National Institute for Health and Care Research Exeter Biomedical Research Centre. The views expressed are those of the author(s) and not necessarily those of the NIHR or the Department of Health and Social Care.

