## Supplementary Figures for "Cell-type-specific DNA methylation dynamics in the prenatal and postnatal human cortex"

**Supplementary Figure 1 - Age estimates derived from an epigenetic clock calibrated on fetal brain are correlated with gestational age in fetal cortex.** Across the 91 fetal cortex samples used in this study there was a strong correlation between actual age and estimated gestational age derived from a fetal brain epigenetic clock (corr = 0.942) [1].

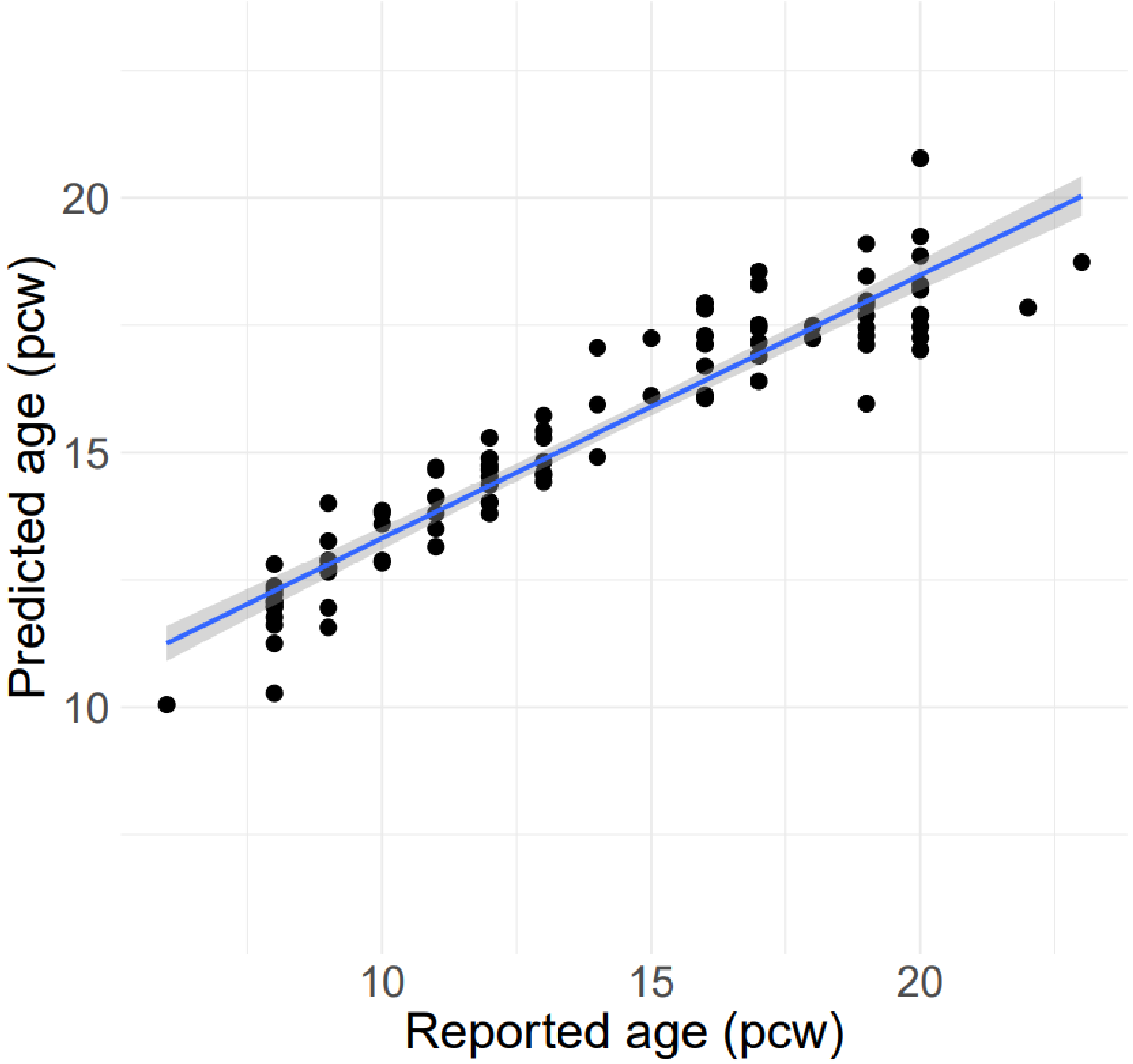

**Supplementary Figure 2 - Developmental changes in DNA methylation in the human cortex are highly correlated with those identified in a previous study of fetal brain across the subset of sites profiled in both studies.** Effect sizes for dDMPs overlapping with sites also profiled in a previous analysis of the fetal brain using the Illumina 450K array (n = 20,502) [2] were highly correlated across studies (corr = 0.824, $p < 1x10^{-320}$).

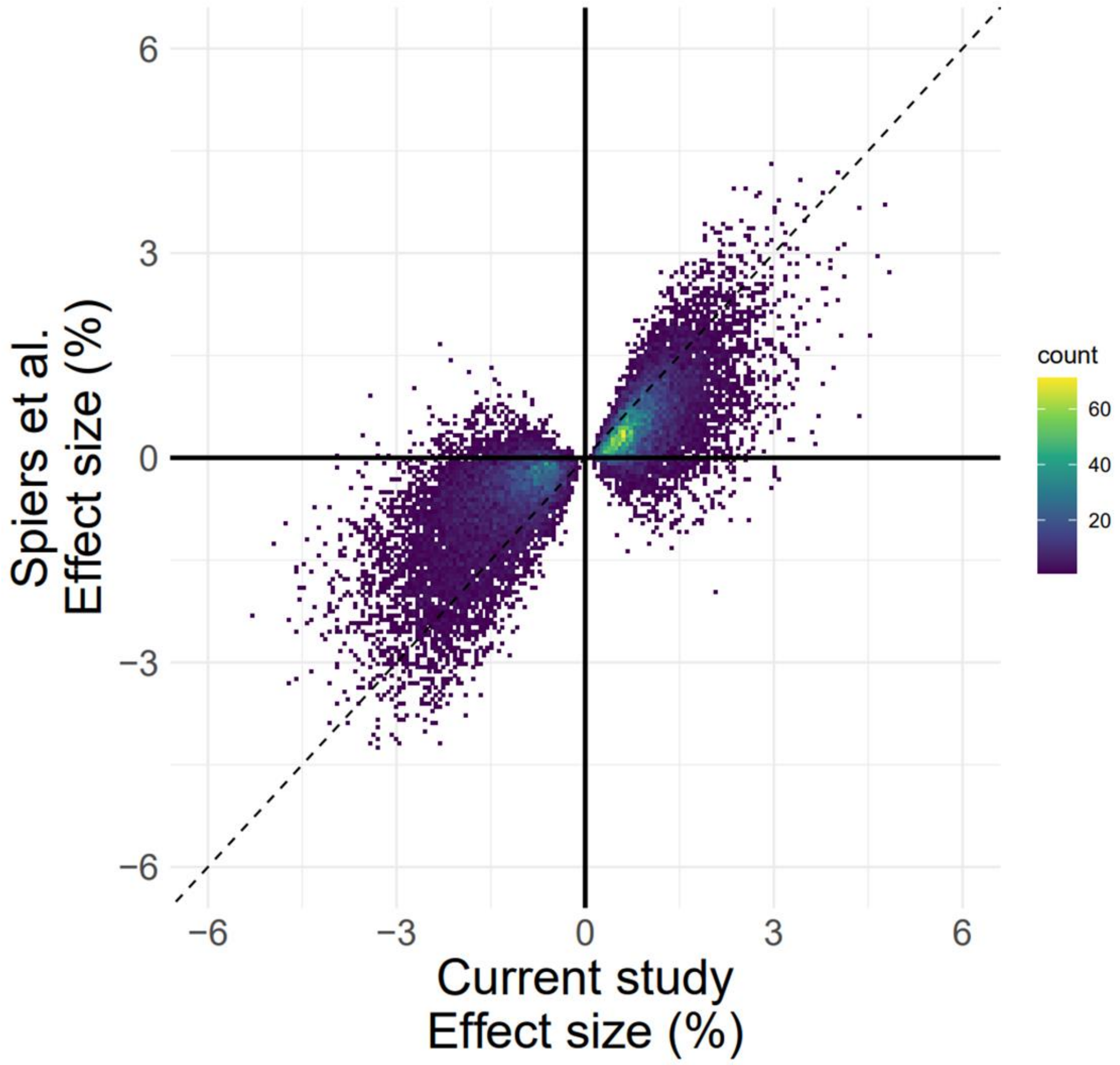

**Supplementary Figure 3 - Distribution of age effect sizes for the 50,913 dDMPs, split by direction of effect.** The mean effect size (absolute change in DNA methylation (%) per week) for hypermethylated dDMPs (red) is smaller than for hypomethylated dDMPs (blue) ($p < 1x10^{-320}$). Dotted lines indicate the mean absolute effect sizes for hypermethylated and hypomethylated dDMPs.

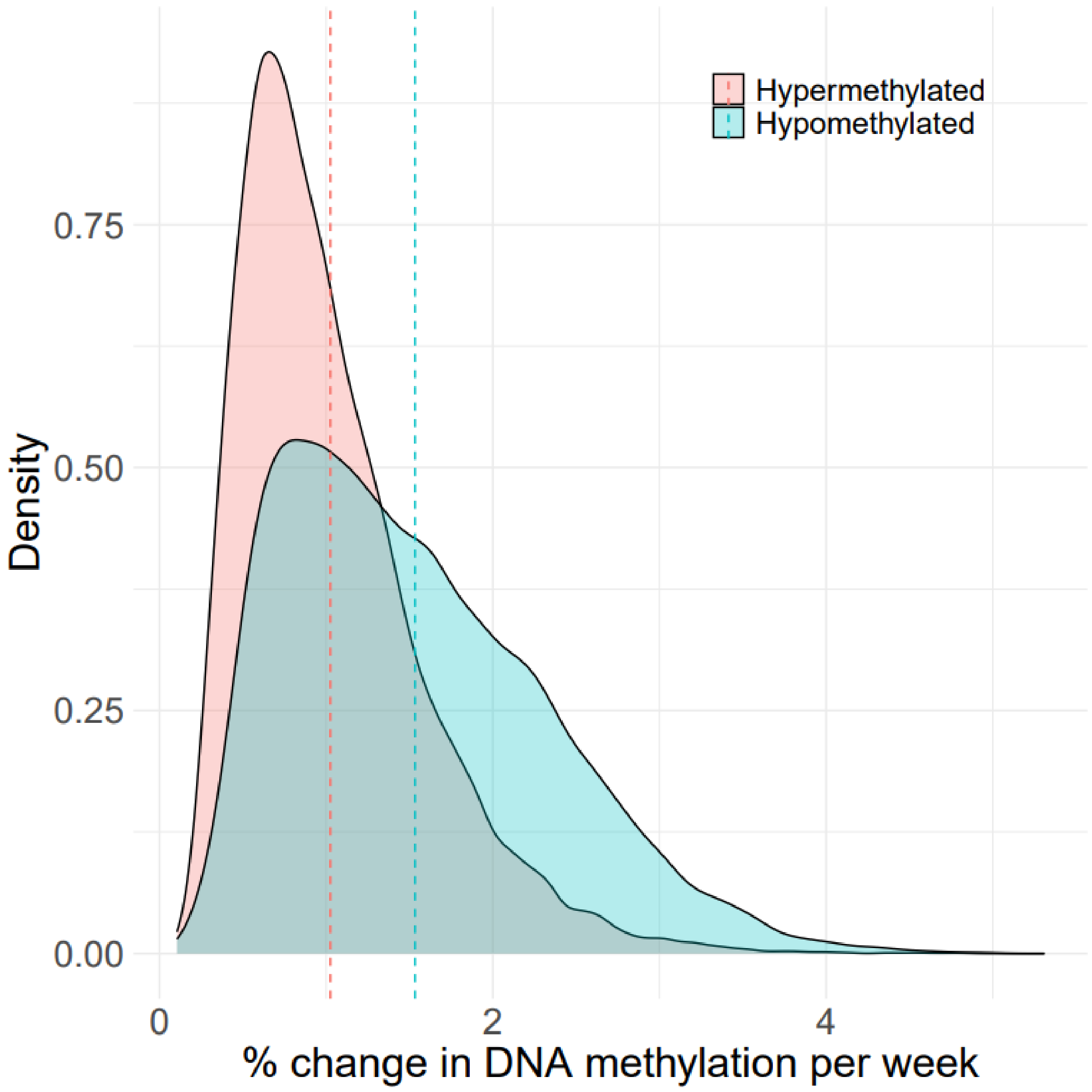

**Supplementary Figure 4 - Mean global DNA methylation decreases across autosomal sites during cortex development. A)** Across all autosomal sites tested (n = 790,180), the mean level of DNA methylation decreases with fetal age (% change in DNA methylation per week = -0.0194, $p = 3.33 \times 10^{-10}$). **B)** The same data is presented as in **A** but the y-axis is scaled from 0 - 100% DNA methylation highlighting the very small magnitude of change across this period. **C)** After excluding autosomal fetal cortex dDMPs, mean global DNA methylation across the remaining 739,839 autosomal sites significantly increases during cortex development (% change in DNA methylation per week = 0.00794, $p = 9.74 \times 10^{-3}$). **D)** The same data is presented as in **C)** but the y-axis is scaled from 0 - 100% DNA methylation.

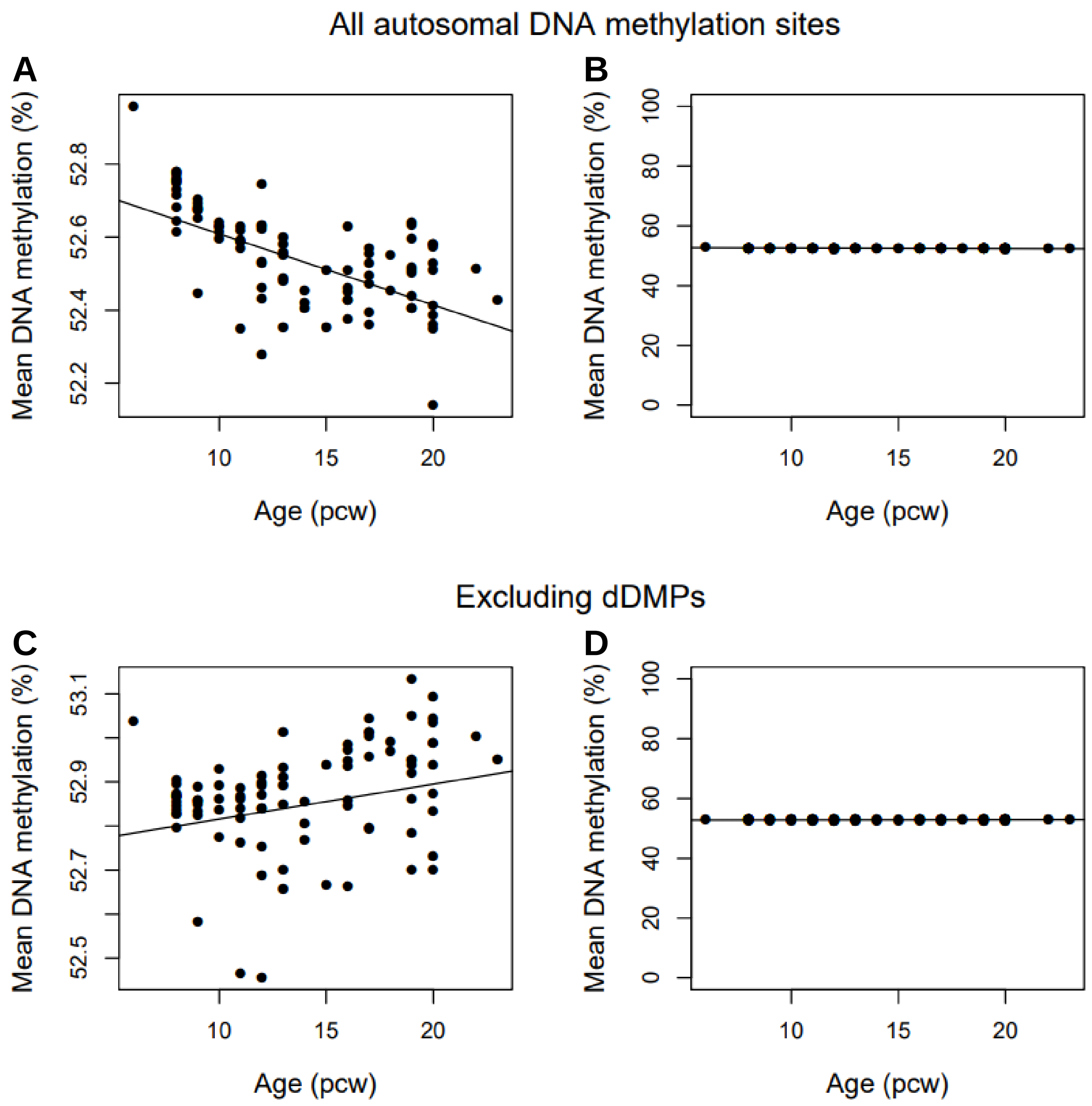

**Supplementary Figure 5 - The distribution of dDMPs across chromosomes was relatively consistent apart from a notable depletion on chromosome 19. A)** Certain chromosomes were characterized by a significant enrichment or depletion of dDMPs (see also **Supplementary Table 4**). Red = significant enrichment of dDMPs, blue = significant depletion of dDMPs, grey = no significant enrichment or depletion.

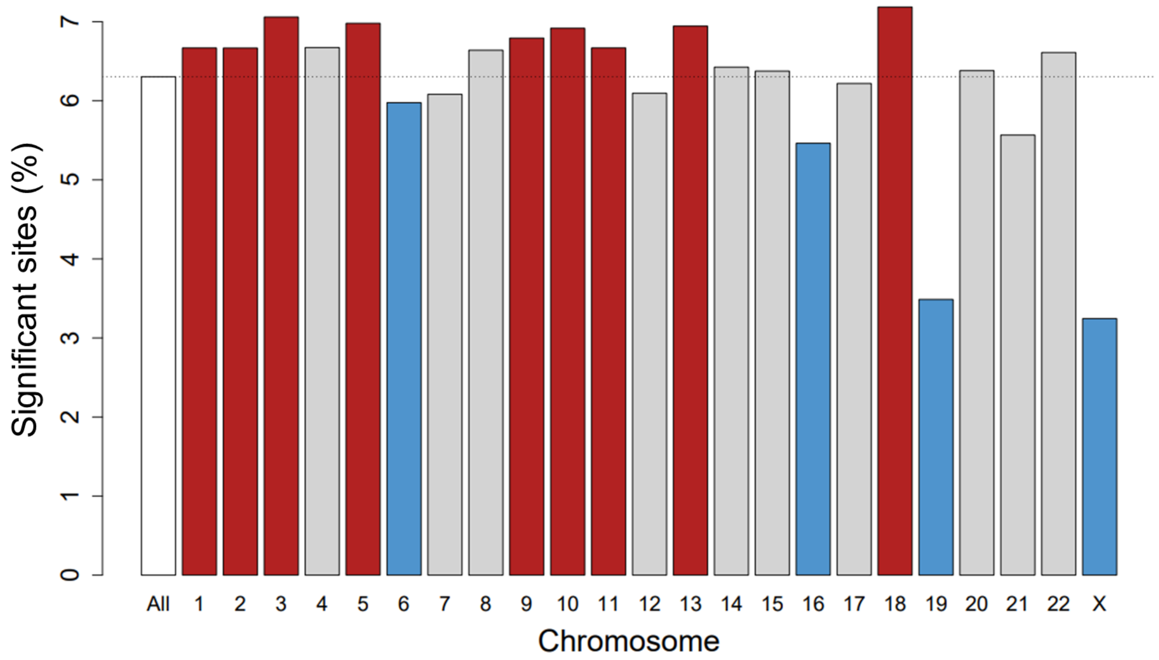

**Supplementary Figure 6 – Cortex dDMPs are enriched in regions of open chromatin identified using scATAC-seq.** Relative effect size from a logistic regression analysis testing for an enrichment of dDMPs within scATAC-seq peaks obtained from a published analysis of 54 human fetal cell-types [3]. 31 out of the 54 cell-types tested were characterized by significant enrichment, with the strongest effect being found for excitatory neurons. The color indicates the direction of effect (red = significant enrichment, blue = significant depletion, gray = non-significant). Full cell-type labels (x-axis) can be found in **Supplementary Table 19**.

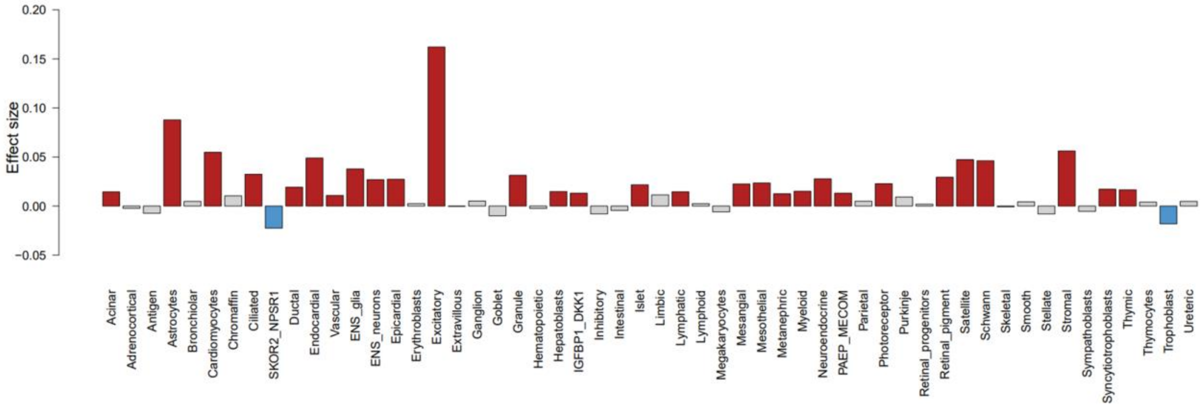

**Supplementary Figure 7 - Age estimates derived from epigenetic clocks calibrated on postnatal samples are strongly correlated with the reported age for the postnatal cortex samples included in this study.** Across the 677 late-fetal and postnatal bulk cortex samples included in this study (26 pcw - 104 years), estimated age was highly correlated to reported age using both **A)** a pan-tissue epigenetic clock (corr = 0.875) [4] and **B)** an epigenetic clock trained on postnatal cortex tissue (corr = 0.941) [5].

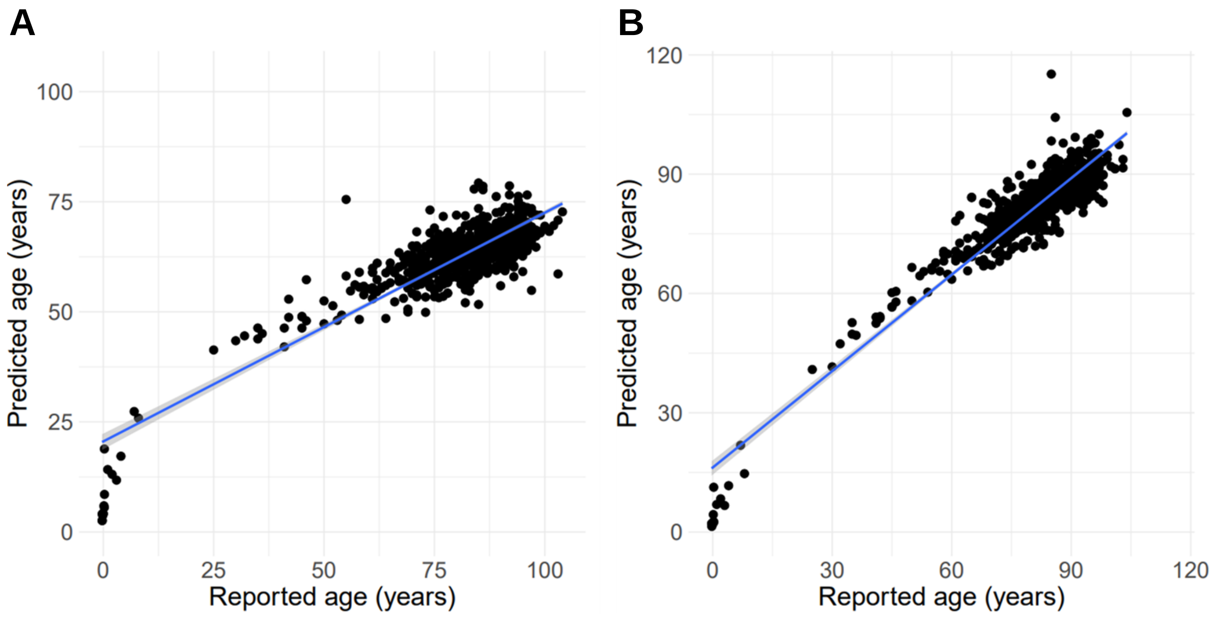

**Supplementary Figure 8 – Life-course trajectories of DNA methylation for the top-ranked hyper- and hypo-methylated dDMPs highlight development-specific effects.** The top hyper- and hypo-methylated dDMPs: cg08125539 (annotated to *IGF2BP1*, postnatal change in DNA methylation (%) per year = -0.0361, p = 0.290) and cg11884704 (annotated to *SLC25A25*, postnatal change in DNA methylation (%) per year = -0.0162, p = 0.493) show little variation in DNA methylation level after birth.

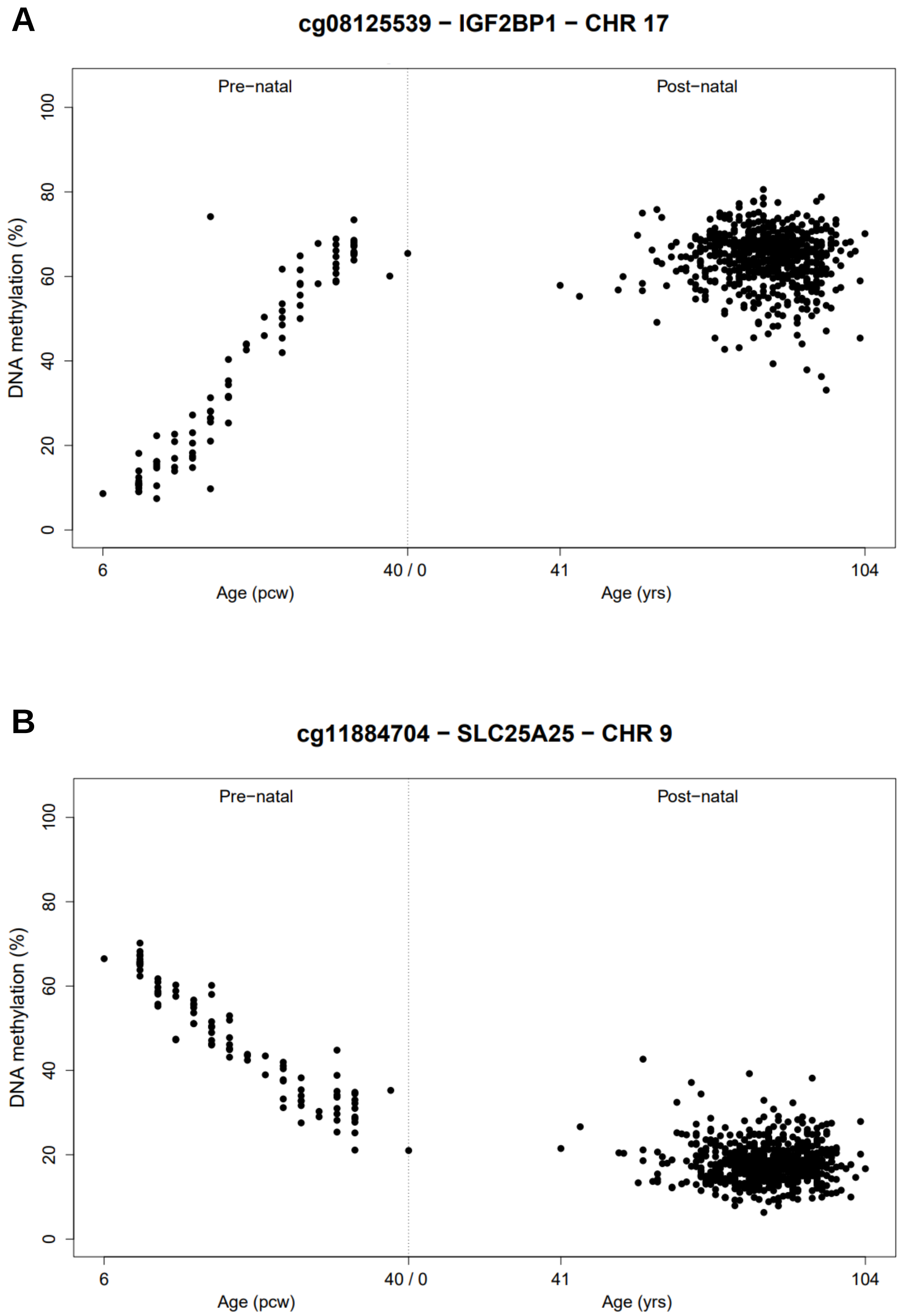

**Supplementary Figure 9 – A large proportion of dDMPs are characterized by nonlinear changes in DNA methylation. A)** Overlap of the 50,913 dDMPs ("Linear dDMPs") identified using a linear regression model and the 73,035 sites classified as showing nonlinear changes in DNA methylation across development using Gaussian process modelling ("Nonlinear sites"). **B)** cg01848383 (annotated to *FOXP1*, a high-confidence SFARI autism gene) is an example of a hypermethylated nonlinear-dDMP and **C)** cg04292453 (annotated to *TRIO*, another high-confidence SFARI autism gene) is an example of a hypomethylated nonlinear-dDMP. A full list of sites at which DNA methylation changes non-linearly across cortex development is given in **Supplementary Table 8**.

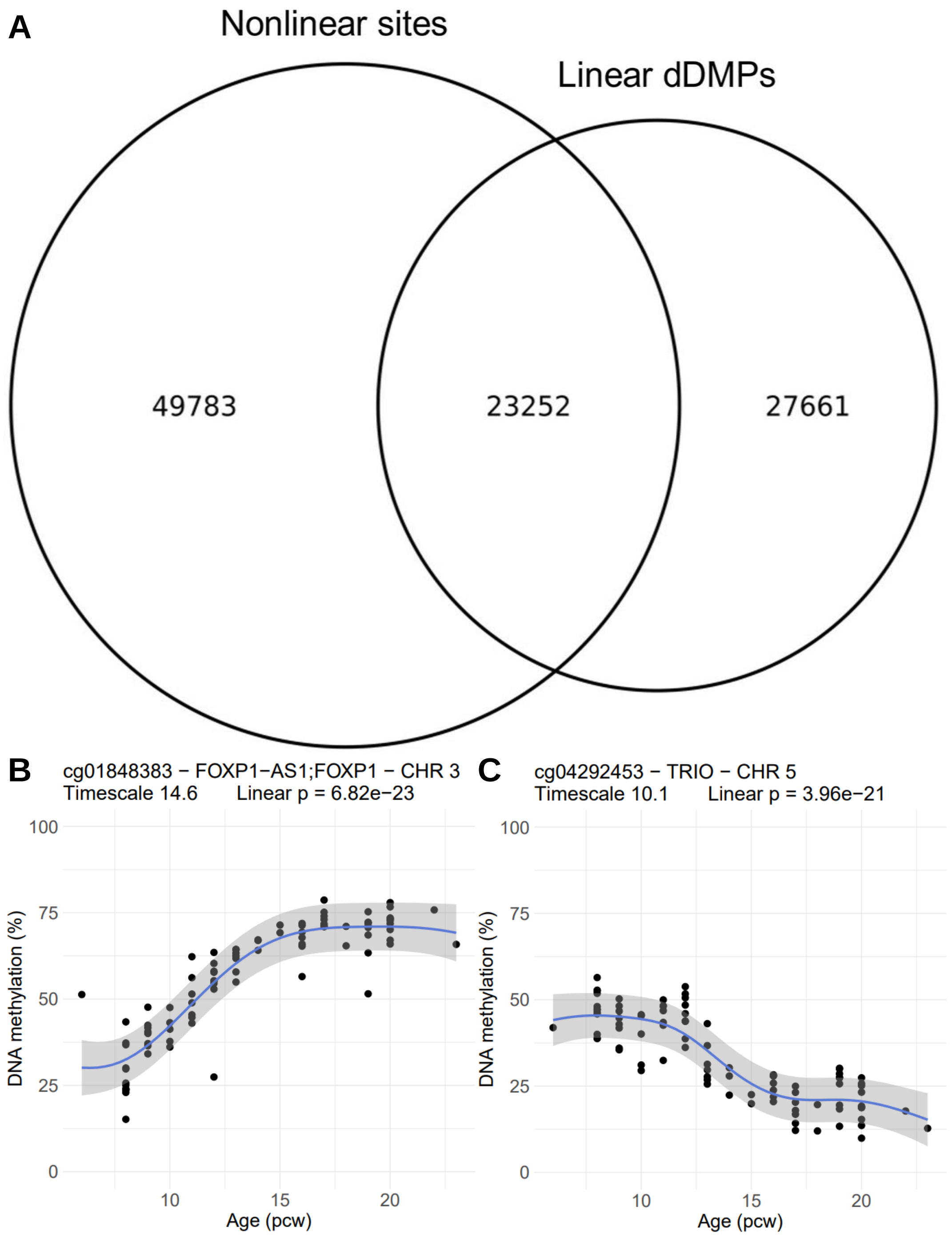

**Supplementary Figure 10 – The expression of *SATB2* is significantly higher than *RBFOX3* (encoding NeuN) during early- and mid-fetal cortex development.** Shown is expression data for *SATB2* and *RBFOX3* from the Brainseq Consortium [6] reported as reads per kilobase per million (RPKM) with standard error shown via shaded area. Developmental stages are defined as follows: Early Midfetal (12 - 18 pcw, n = 23), Midfetal (18 - 19 pcw, n = 14), Late Midfetal (19 - 26 pcw, n = 13), Late Fetal (26 - 40 pcw, n = 3), Early Infancy (0 - 180 days, n = 16), Early Childhood (1 - 6 years, n = 12), Late Childhood (6 - 13 years, n = 4), Adolescence (13 - 20 years, n = 47), Young Adulthood (20 - 30 years, n = 35), Mid Adulthood (30 - 60 years, n = 139), Late Adulthood (≥ 60 years, n = 28). Expression of *RBFOX3* was significantly lower than expression of *SATB2* in prenatal cortex ($t = -14.562$, $p < 2.2e-16$) but higher than expression of *SATB2* in postnatal cortex ($t = 25.01$, $p < 2.2e-16$).

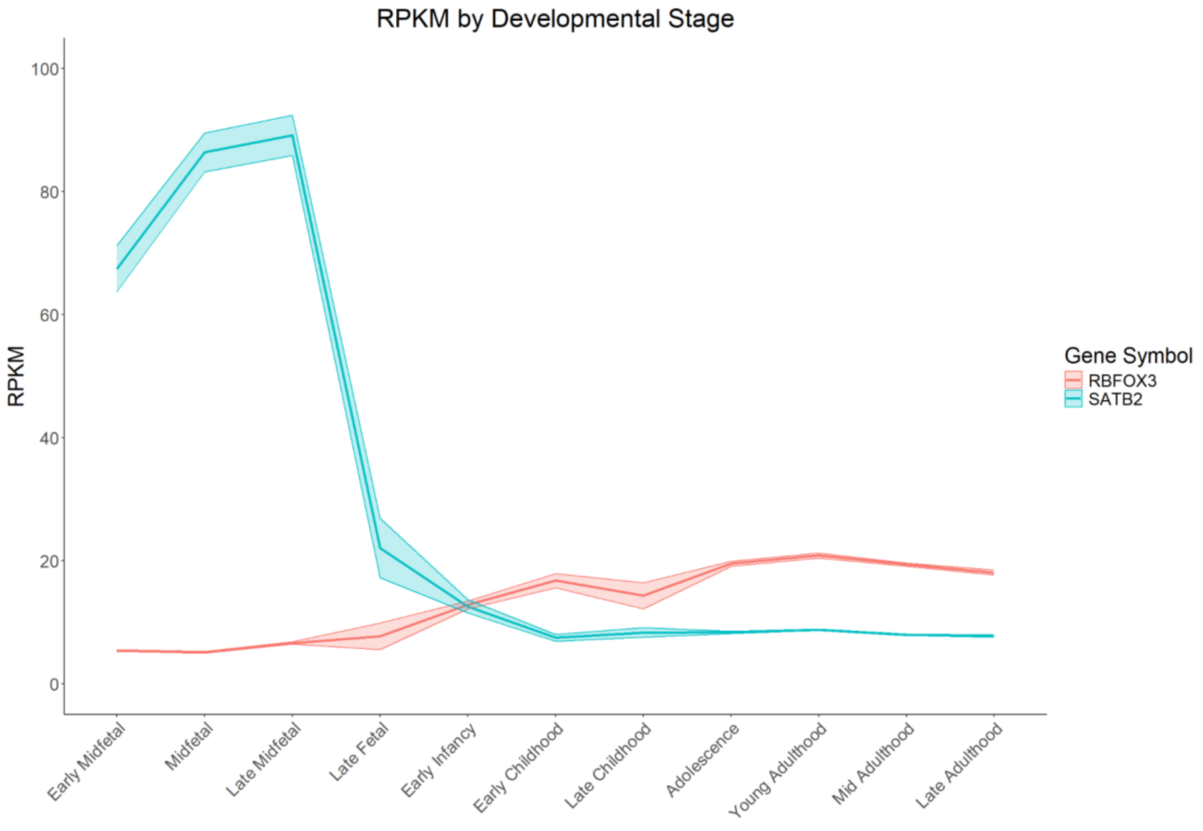

**Supplementary Figure 11 – Broad neuronal co-expression of *RBFOX3* and *SATB2* in postnatal cortex neurons.** Shown for total nuclei (bulk cortex) and FANS-isolated SATB2+ and NeuN+ nuclei is the expression of *SATB2* and *RBFOX3* across annotated cell-types quantified using single nuclei RNA-seq.

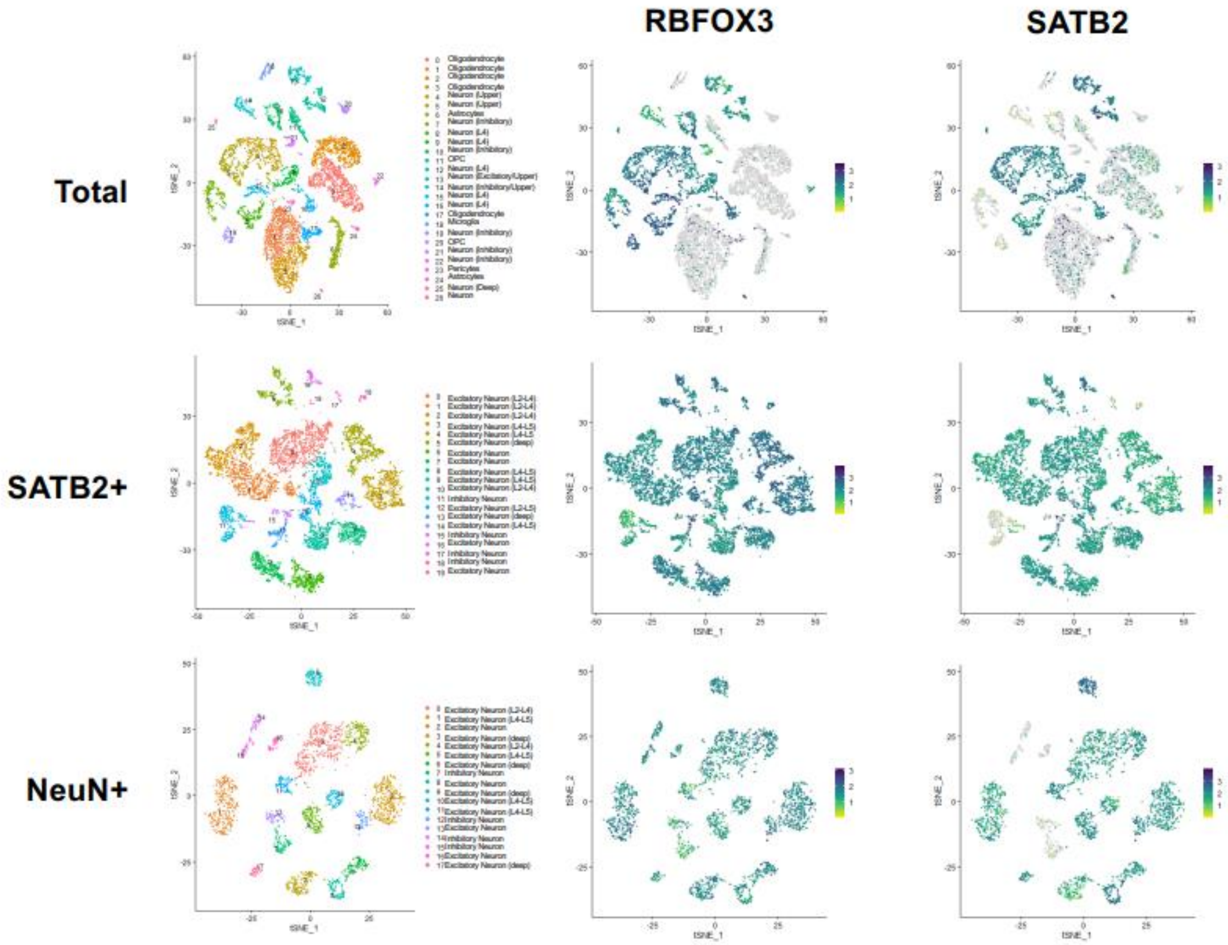

**Supplementary Figure 12 – *SATB2* expression is higher than *RBFOX3* expression in nuclei isolated from fetal cortex.** Shown for total nuclei (bulk cortex) and FANS-isolated SATB2+ nuclei is the expression of *SATB2* and *RBFOX3* across annotated cell-types quantified using snRNA-seq. The higher expression of *SATB2* in the fetal cortex parallels the data shown in S**upplementary Figure 11**.

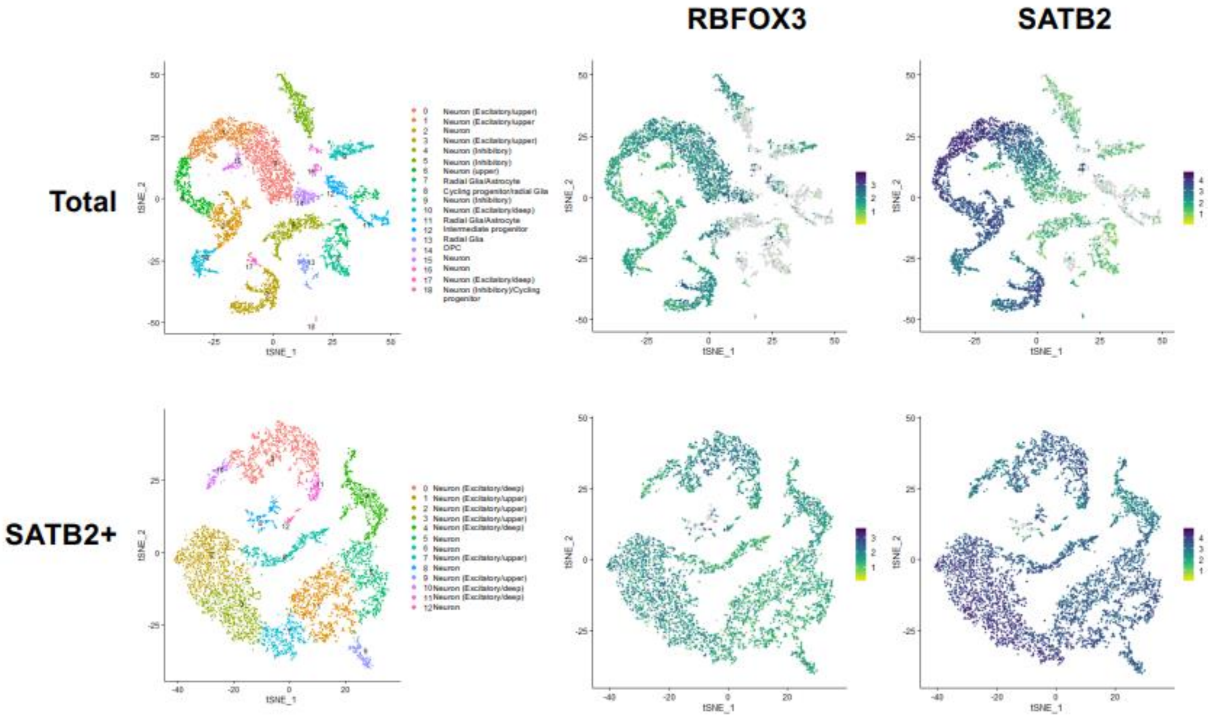

**Supplementary Figure 13 – A comparison of DNA methylation differences between neuronal and non-neuronal nuclei isolated from fetal and adult cortex.** Neuronal vs non-neuronal DNA methylation differences in adult (n = 212 donors, x-axis) and fetal (n = 37 donors, y-axis) cortex for **A)** all sites tested in both (n = 693,895), **B)** sites identified as showing significant neuronal vs non-neuronal differences in fetal cortex (n = 6,531) and **C)** sites identified as showing significant neuronal vs non-neuronal differences in adult cortex (n = 453,675). Effect size represents the difference in mean DNA methylation (%) between neuronal and non-neuronal nuclei populations. 'Count' in **A** represents the number of DNA methylation sites per bin.

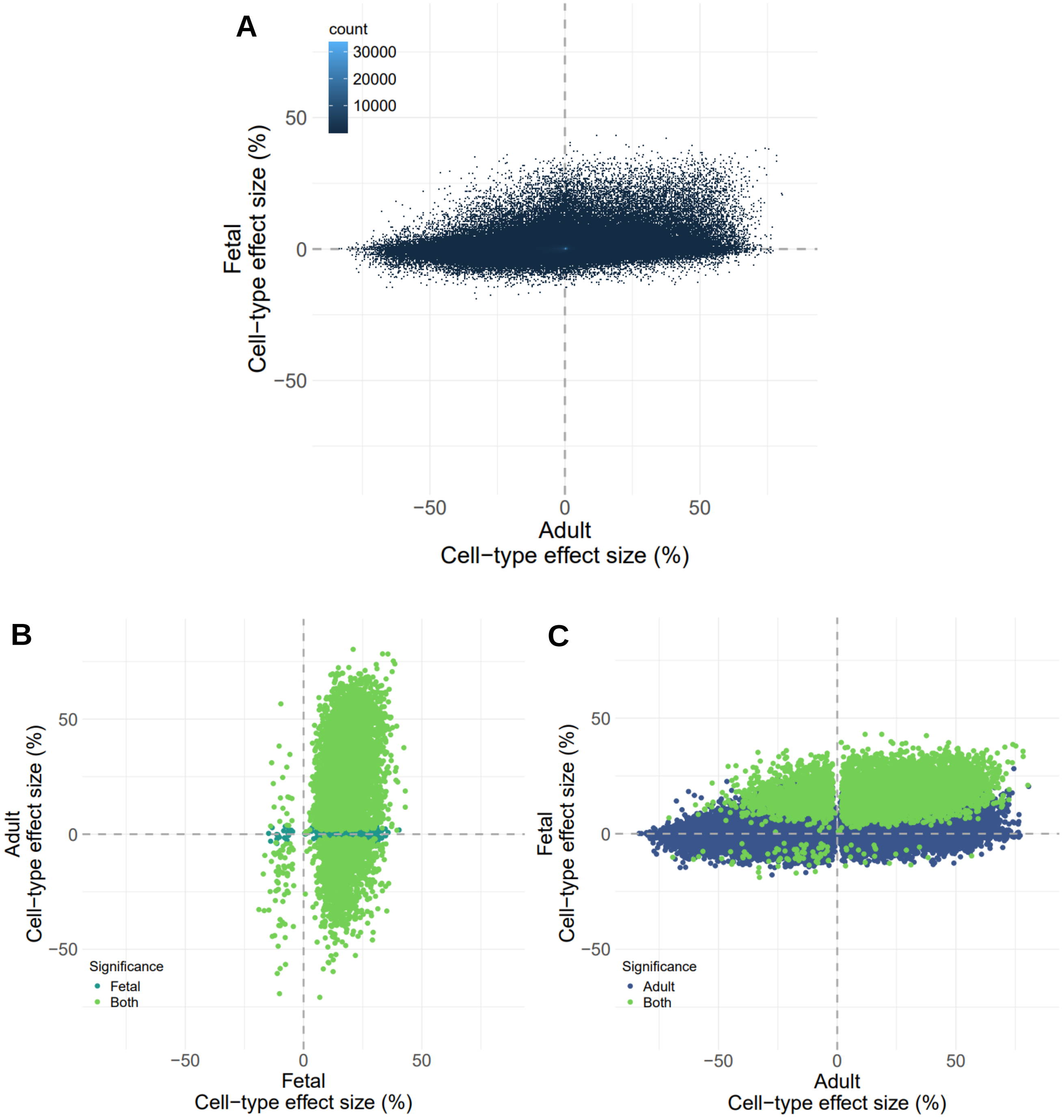

**Supplementary Figure 14 – Comparison of developmental changes in DNA methylation identified in bulk cortex and FANS-isolated nuclei populations from neuronal and non-neuronal cells.** Shown for the 42,114 bulk cortex dDMPs also tested in FANS-isolated populations is the correlation of developmental changes between **A)** bulk fetal cortex and neuronal nuclei, **B)** bulk cortex and non-neuronal nuclei and **C)** neuronal and non-neuronal nuclei. Effect size represents the percentage change in DNA methylation per post-conception week.

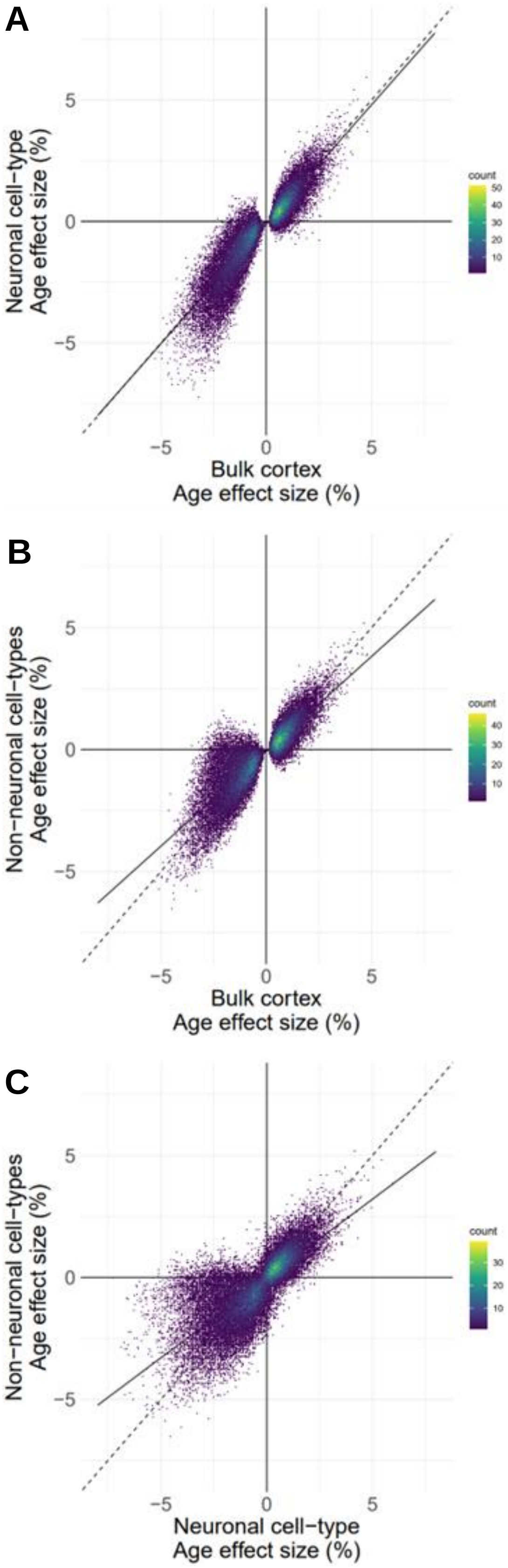

**Supplementary Figure 15 – Overlap of developmental changes in DNA methylation identified in neuronal and non-neuronal nuclei isolated from the fetal cortex.** Of the 1,872 neuronal and 820 non-neuronal dDMPs, 272 are consistent between cell-types ($p < 9x10^{-8}$), all of which are concordant in direction.

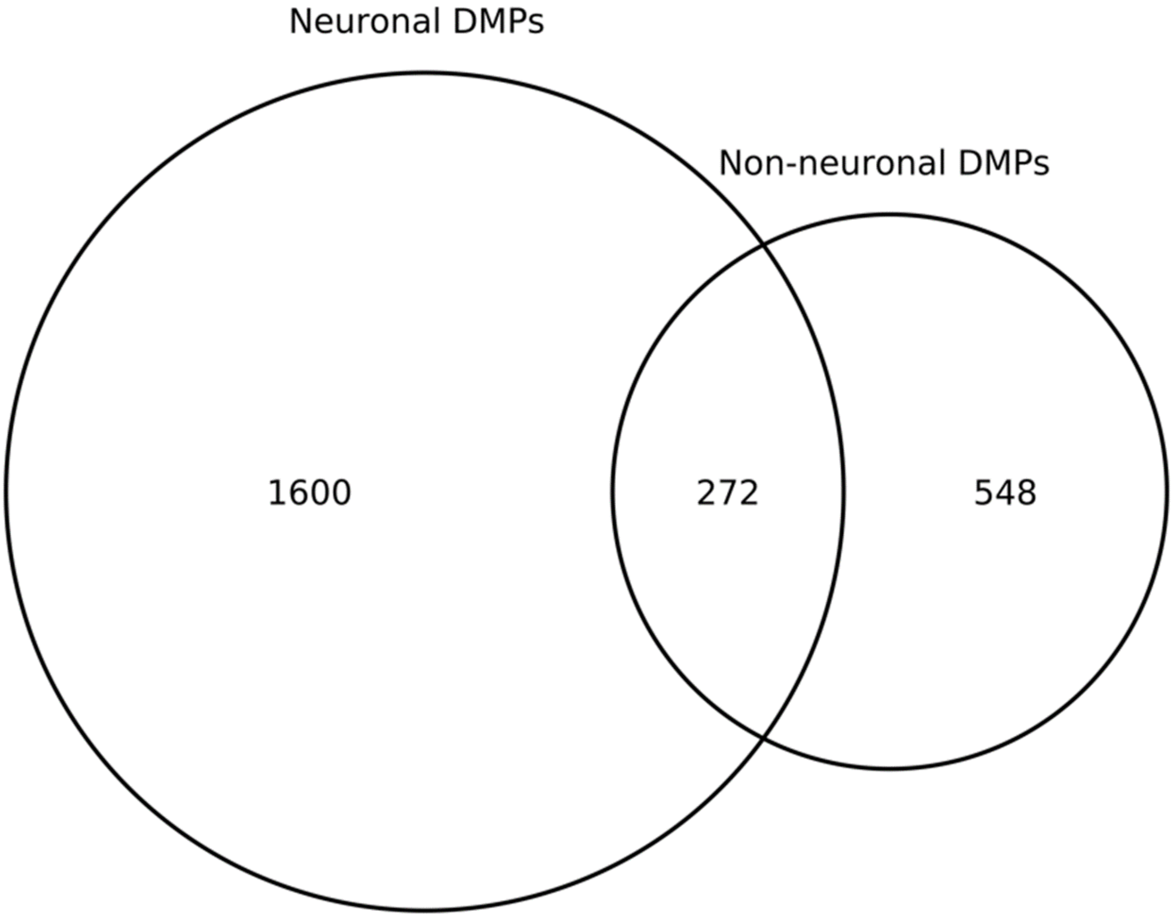

**Supplementary Figure 16 – Venn diagram showing the overlap of dDMPs identified in bulk fetal cortex and purified nuclei populations.** Overlap of the 42,114 bulk cortex dDMPs ($p < 9x10^{-8}$) also tested in our FANS-isolated samples with the 4,088 neuronal dDMPs ($p < 1.19x10^{-6}$) and 1,820 non-neuronal dDMPs ($p < 1.19x10^{-6}$).

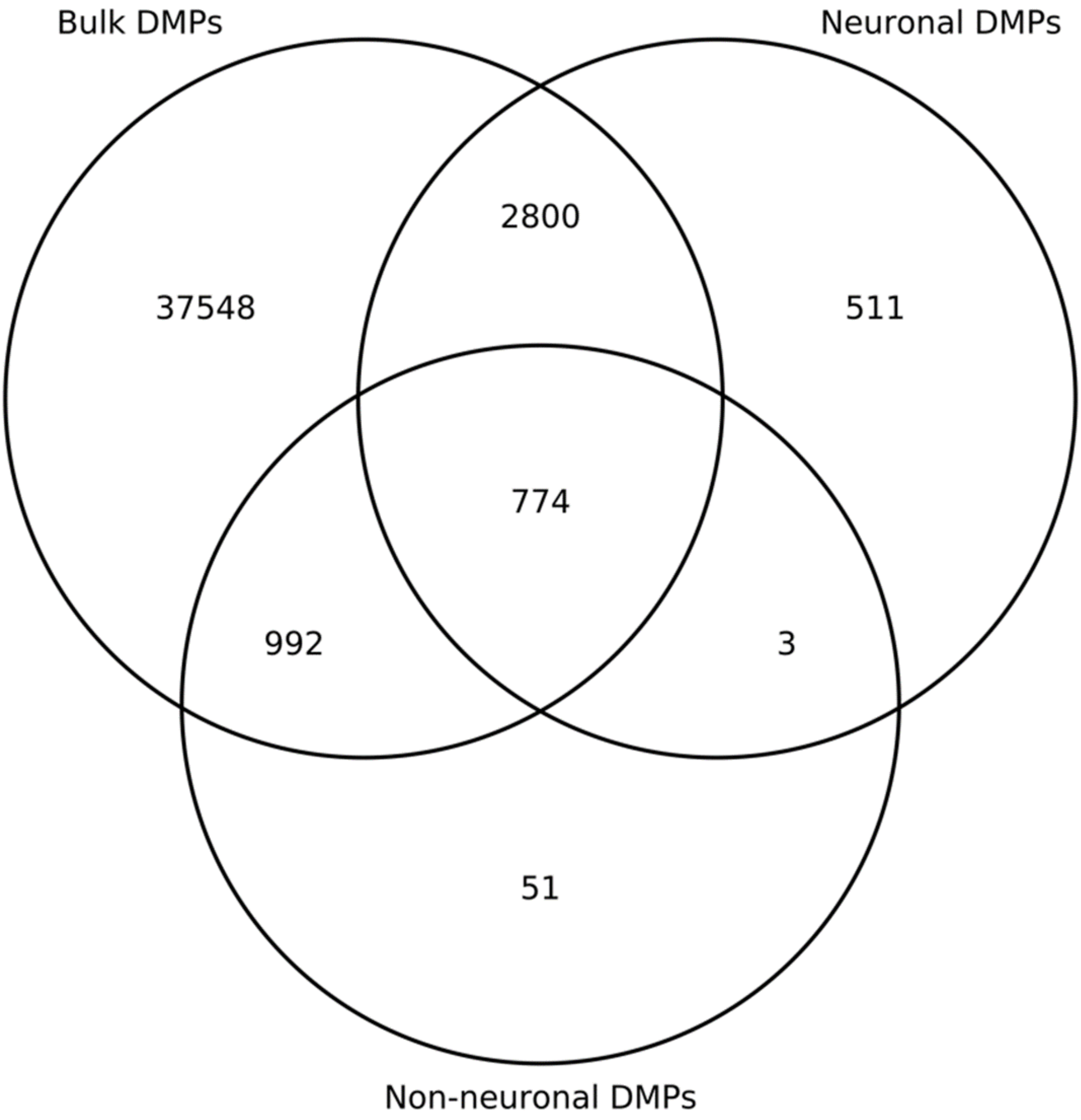

**Supplementary Figure 17 – Many developmental changes in DNA methylation identified the bulk fetal cortex reflect changes that are specific to neuronal or non-neuronal cells.** Shown are examples of bulk cortex dDMPs that are characterized by **A)** neuronal-specific changes, **B)** non-neuronal-specific changes and **C-D)** opposite direction changes between the neuronal and non-neuronal populations.

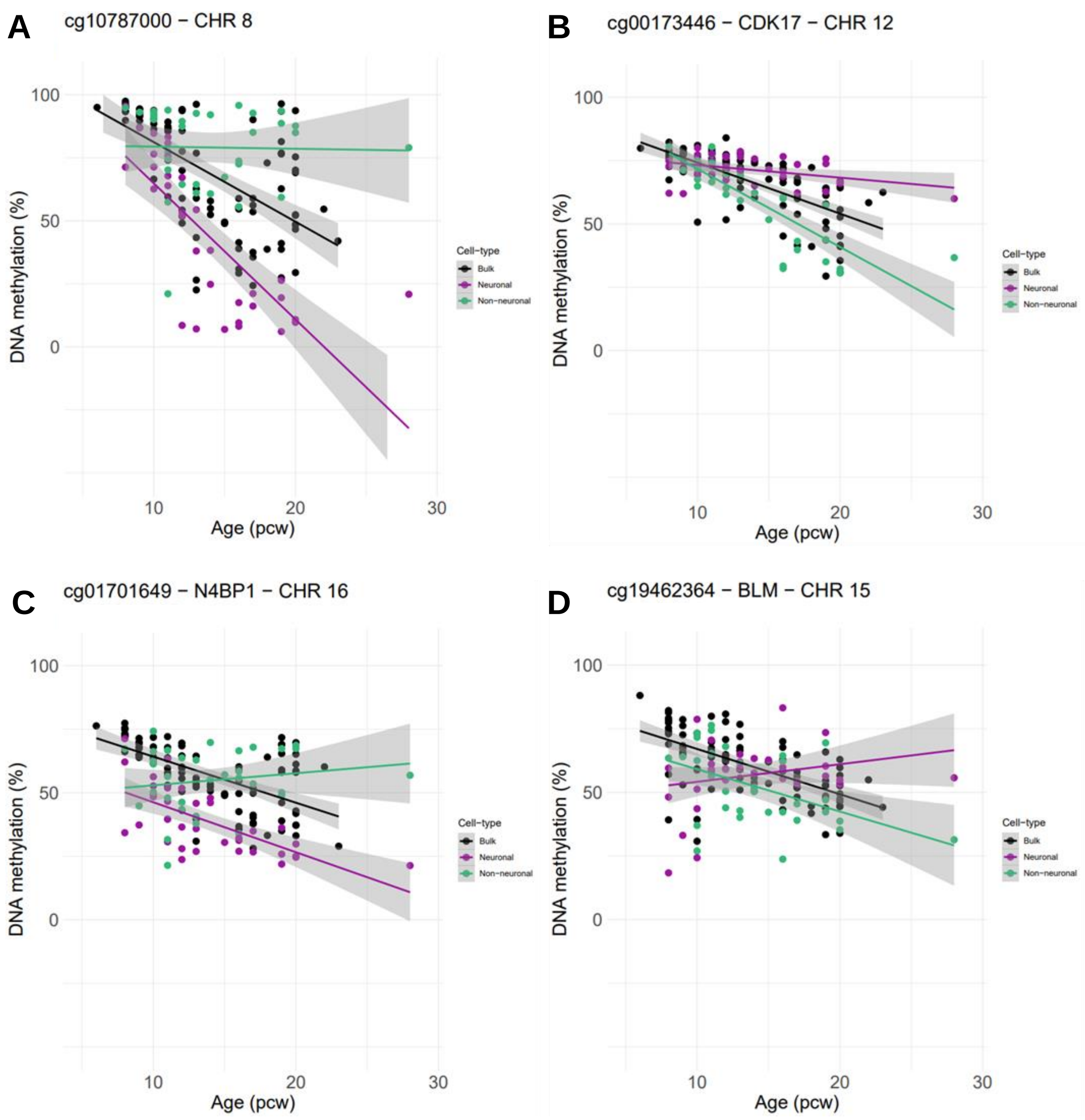

**Supplementary Figure 18 – Identification of cell-type specific changes in DNA methylation in human cortex development.** Comparison of age effect sizes for **A)** dDMPs identified in either neuronal or non-neuronal nuclei (n = 2,420, corr = 0.564), **B)** neuronal dDMPs (n = 1,872, corr = 0.634) and **C)** non-neuronal dDMPs (n = 820, corr = 0.873). Effect size represents the change in DNA methylation (%) per post-conception week. The results show that developmental changes identified in non-neuronal nuclei are broadly reflected in neuronal nuclei, but a large proportion of neuronal dDMPs are cell-type-specific.

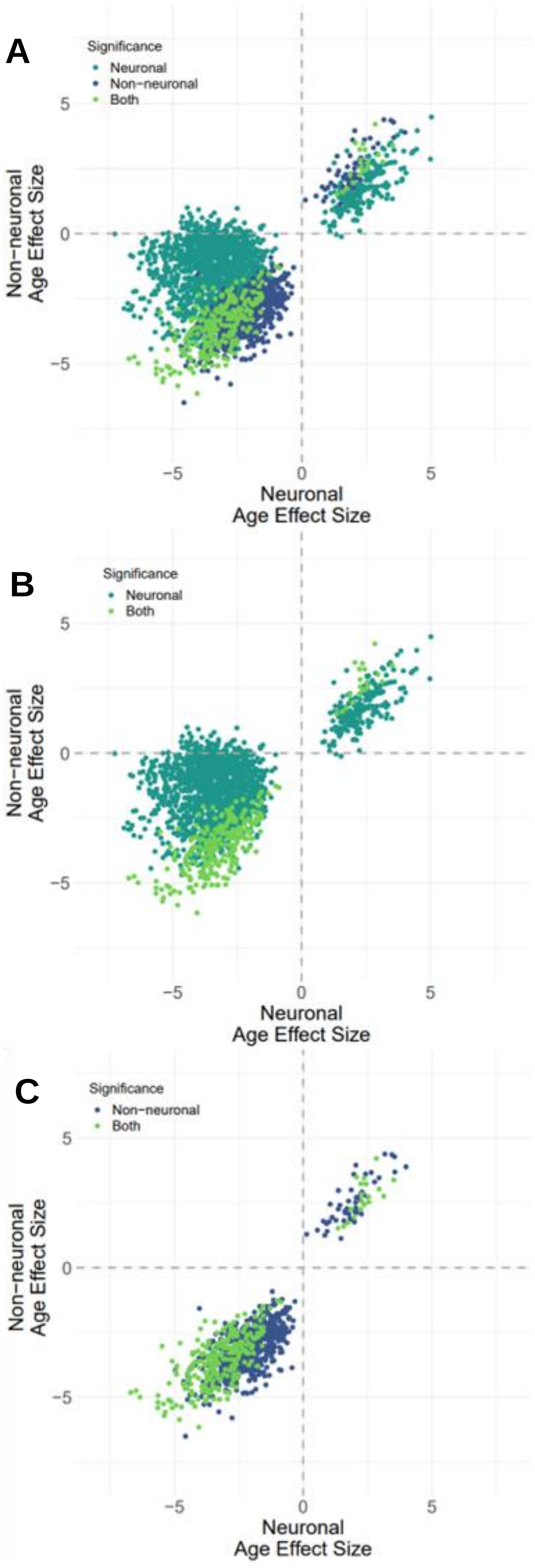

**Supplementary Figure 19 – Examples of sites characterized by cell-type-specific developmental changes in DNA methylation in the fetal cortex. A)** cg12430457 (annotated to *SYT1*, a gene involved in synaptic vesicle exocytosis [7]) becomes developmentally hypomethylated in neuronal nuclei but not non-neuronal nuclei. **B)** cg09165170 (annotated to *PEX14*, a gene that encodes a peroxisomal protein crucial for import of cargo into peroxisomes which are critical in oligodendrocytes for the maintenance of myelination [8]) becomes developmentally hypermethylated in neuronal nuclei but not non-neuronal nuclei. **C)** cg13609939 (annotated to *OAT*, a gene involved in the glutamate metabolic pathway [9,10]) shows a greater rate of hypomethylation in non-neuronal nuclei compared to neuronal nuclei. **D)** cg23699648 (annotated to *GJA1*, encoding a connexin protein that plays an important role in neural development [11]) shows a greater rate of hypermethylation in neuronal nuclei compared to non-neuronal nuclei. A full list of sites showing cell-type-specific developmental changes in DNA methylation in the fetal cortex is given in **Supplementary Table 11**.

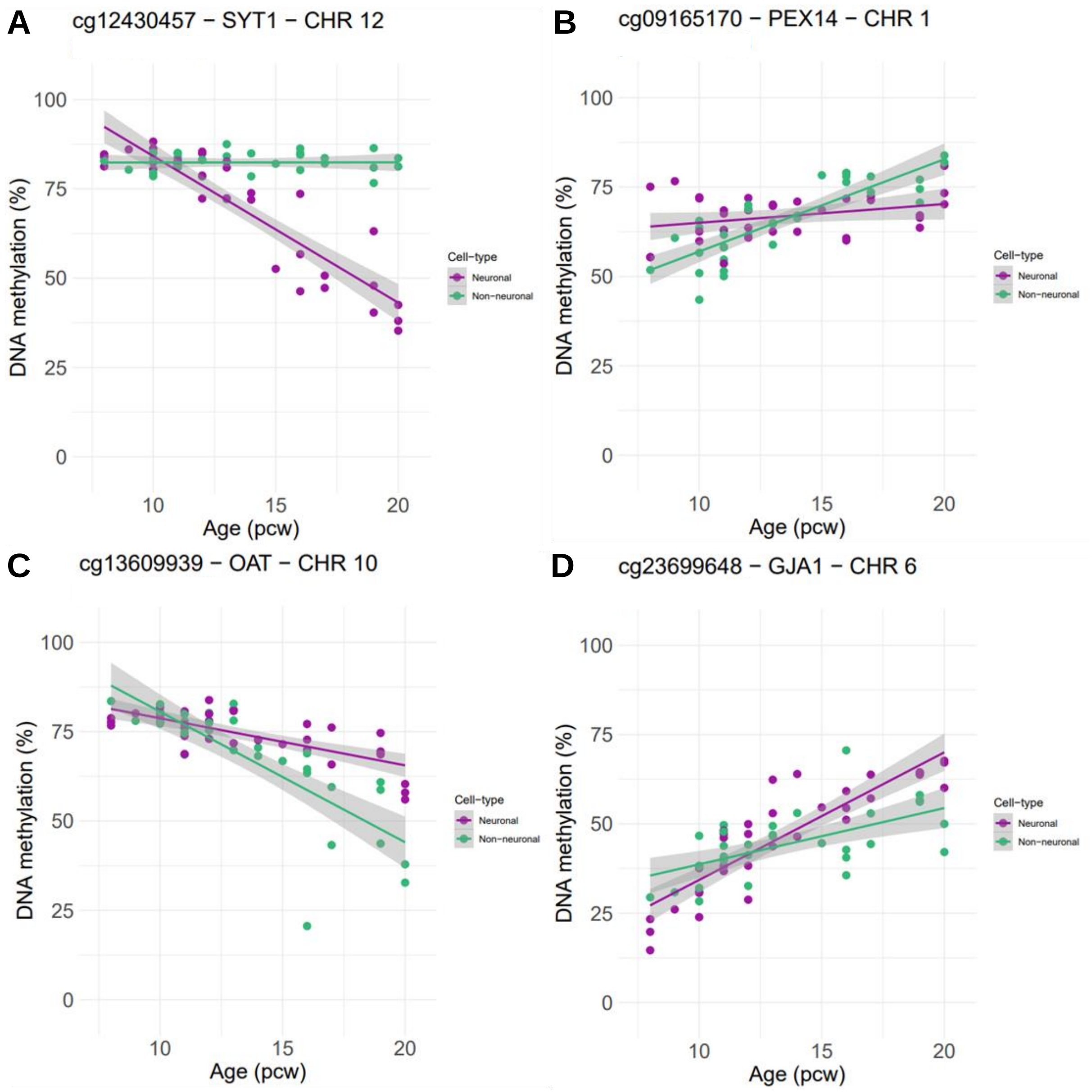

**Supplementary Figure 20 – Enrichment of cell-type-specific dDMPs in regions of open chromatin identified in relevant cell-types by scATAC-seq.** Relative effect size from a logistic regression analysis testing for an enrichment of dDMPs within scATAC-seq peaks obtained from a published analysis of 54 human fetal cell-types [3]. **A)** Enrichment of neuron-specific autosomal dDMPs (n = 1,596) within cell-type-specific ATAC-seq peaks. Excitatory neurons demonstrate the single clearest enrichment, followed by astrocytes and stromal cells (see also **Supplementary Table 13**). **B)** Enrichment of non-neuron-specific autosomal dDMPs (n=548) within cell-type-specific ATAC-seq peaks. There is a clear enrichment of non-neuronal brain cell-types including astrocytes and Schwann cells (see also **Supplementary Table 14**).

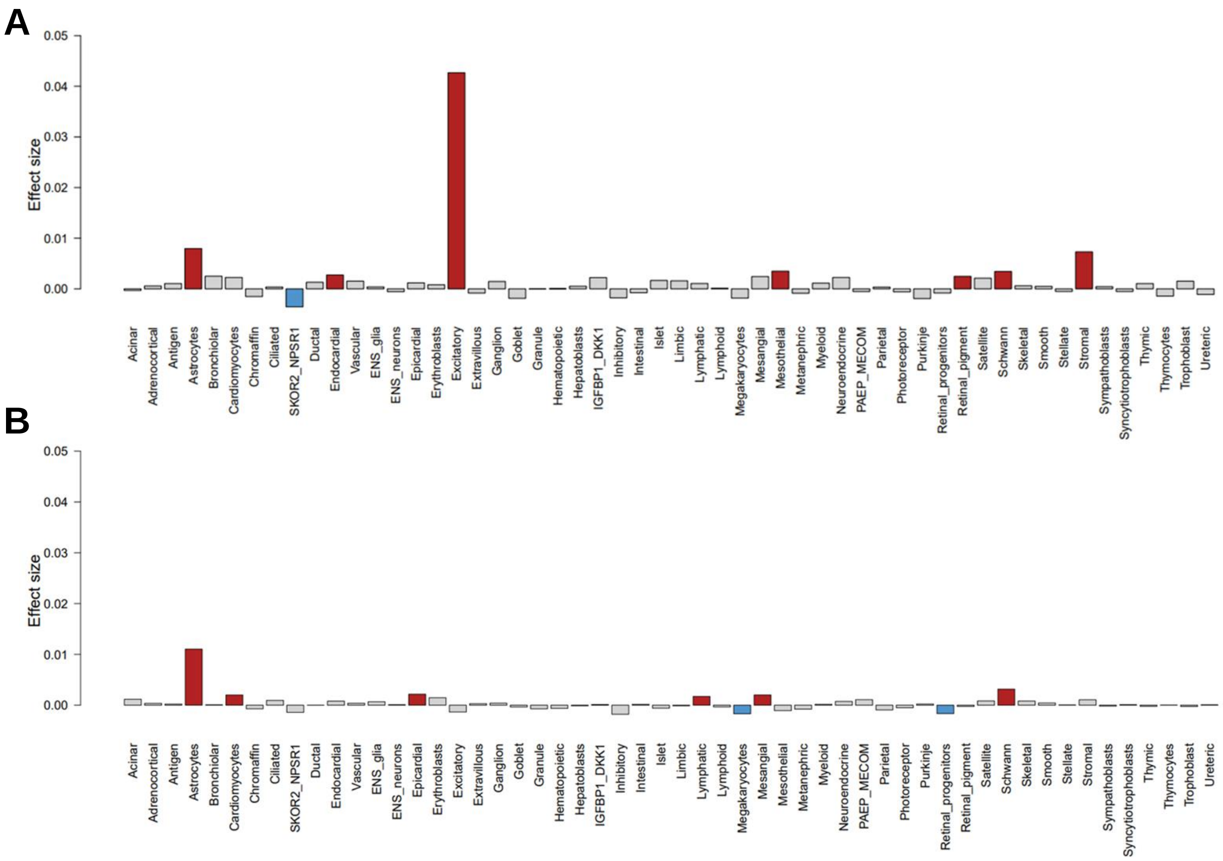

**Supplementary Figure 21 – Proportion of bulk fetal cortex dDMPs annotated to autism (SFARI) and schizophrenia (SCHEMA) genes. A)** Percentage of genes that contain at least one dDMP across four gene lists: i) all 26,728 genes annotated to sites on the array (gray), ii) the 233 SFARI genes (blue), iii) the 32 SCHEMA genes (green) and iv) the 259 combined unique SFARI and SCHEMA genes (purple). **B)** Percentage of all sites annotated to each of the gene lists that is a dDMP.

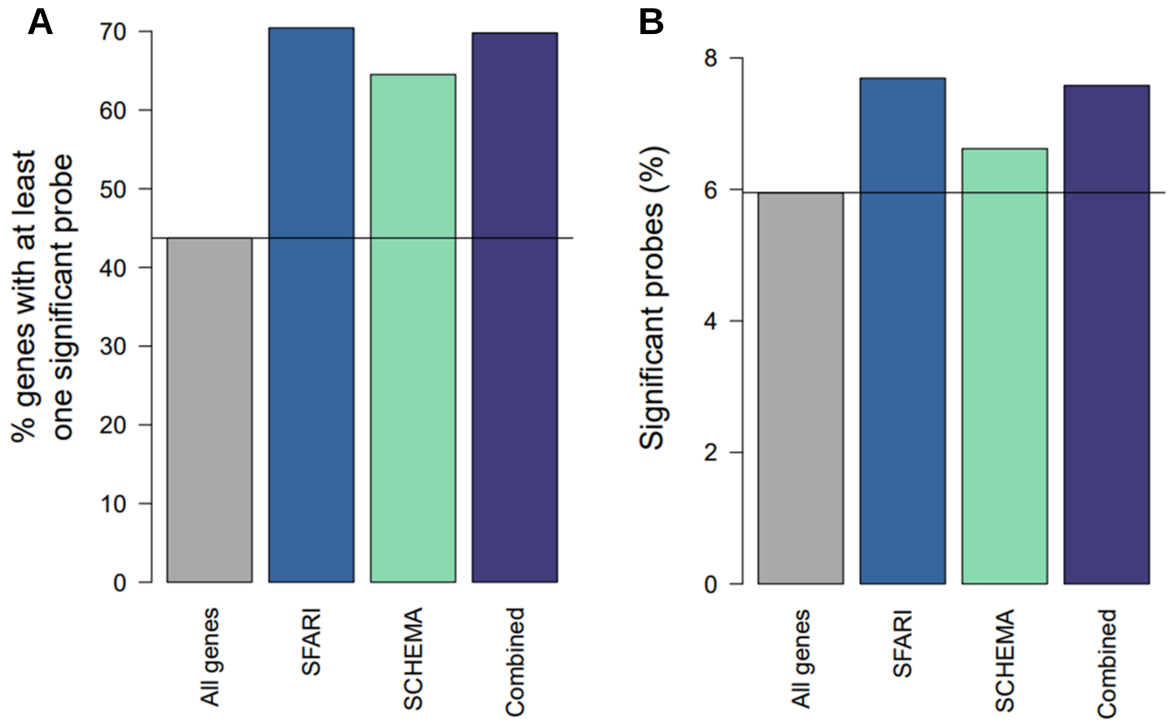

**Supplementary Figure 22 – Proportion of neuronal and non-neuronal dDMPs annotated to autism (SFARI) and schizophrenia (SCHEMA) genes. A)** Percentage of genes that contain at least one neuronal dDMP across four gene lists: i) all 26,455 genes annotated to sites on the array (grey), ii) the 233 SFARI genes (blue), and iii) the 32 SCHEMA genes (green) and iv) the 259 combined SFARI and SCHEMA genes (purple). **B)** Percentage of all sites annotated to each gene list that is a neuronal dDMP. **C)** Percentage of genes that contain at least one non-neuronal dDMP across each of the four gene lists. **D)** Percentage of all sites annotated to each gene list that is a non-neuronal dDMP.

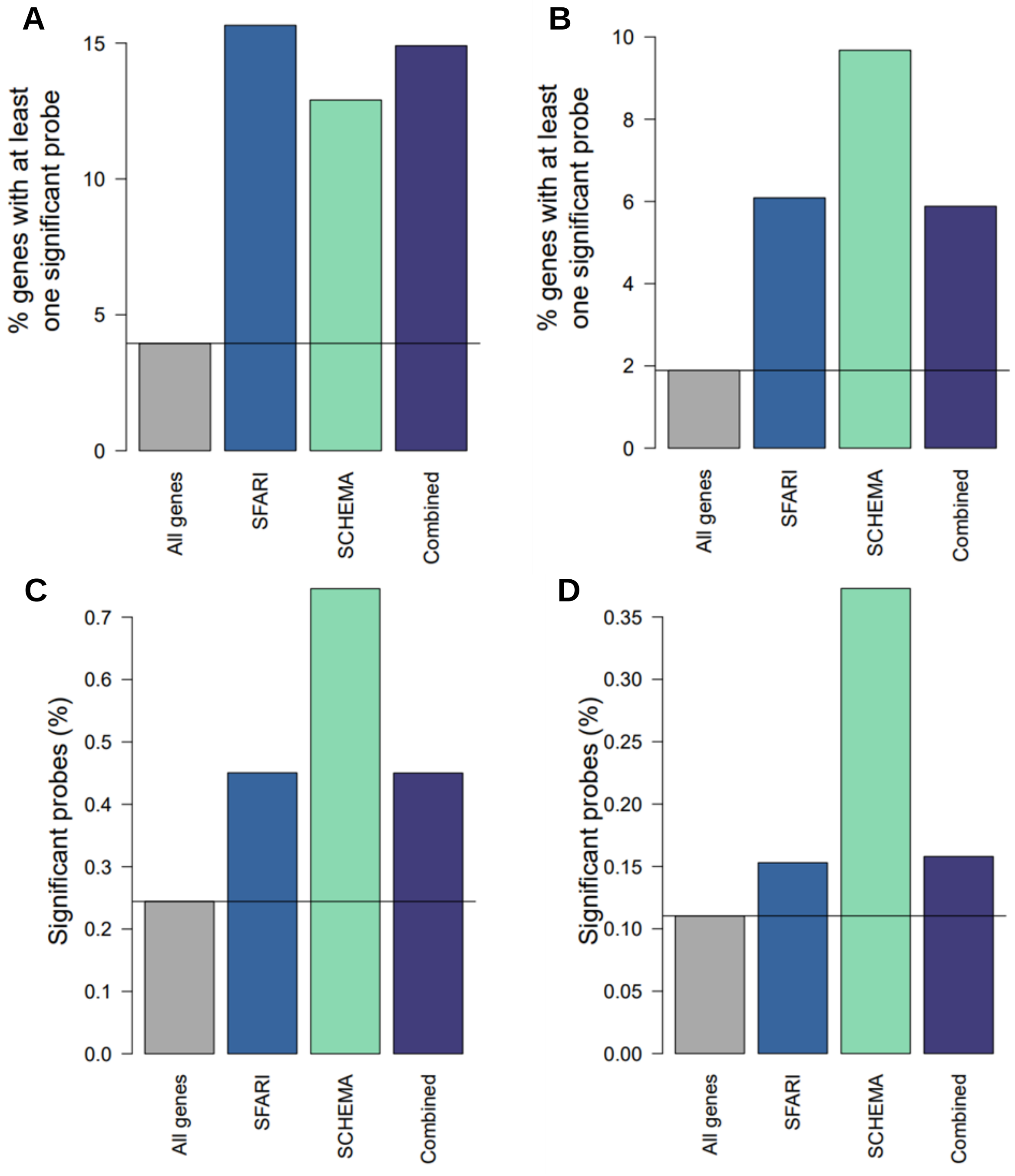

**Supplementary Figure 23 – MAGMA gene set enrichment.** Enrichment of genes annotated to bulk cortex, neuronal and non-neuronal dDMPs for common variants associated with **A)** autism and **B)** schizophrenia using MAGMA [12]. Circles indicate enrichment effect size +/- 95% confidence interval. p = unadjusted p-value.

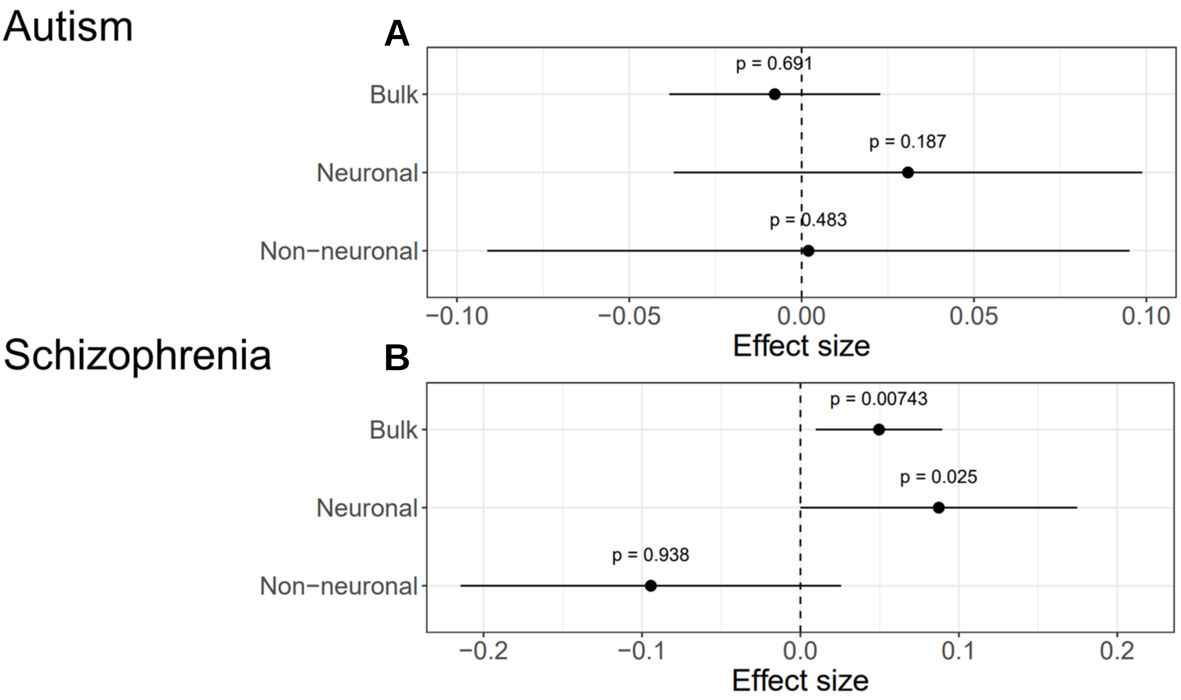

**Supplementary Figure 24 – Log-likelihood ratio to refine nonlinear sites.** Shown is timescale against the log-likelihood ratios (LLRs) of **A)** the Matern 5/2 (nonlinear) vs constant kernel and **B)** Matern 5/2 vs linear kernel, for the 175,419 nonlinear sites identified from the Gaussian process model as having the greatest marginal log-likelihood for the Matern 5/2 kernel. Sites with LLR < 2 and timescale < 10 were excluded, leaving 73,638 high-confidence nonlinear sites with biologically meaningful timescales between 10.0 and 106.

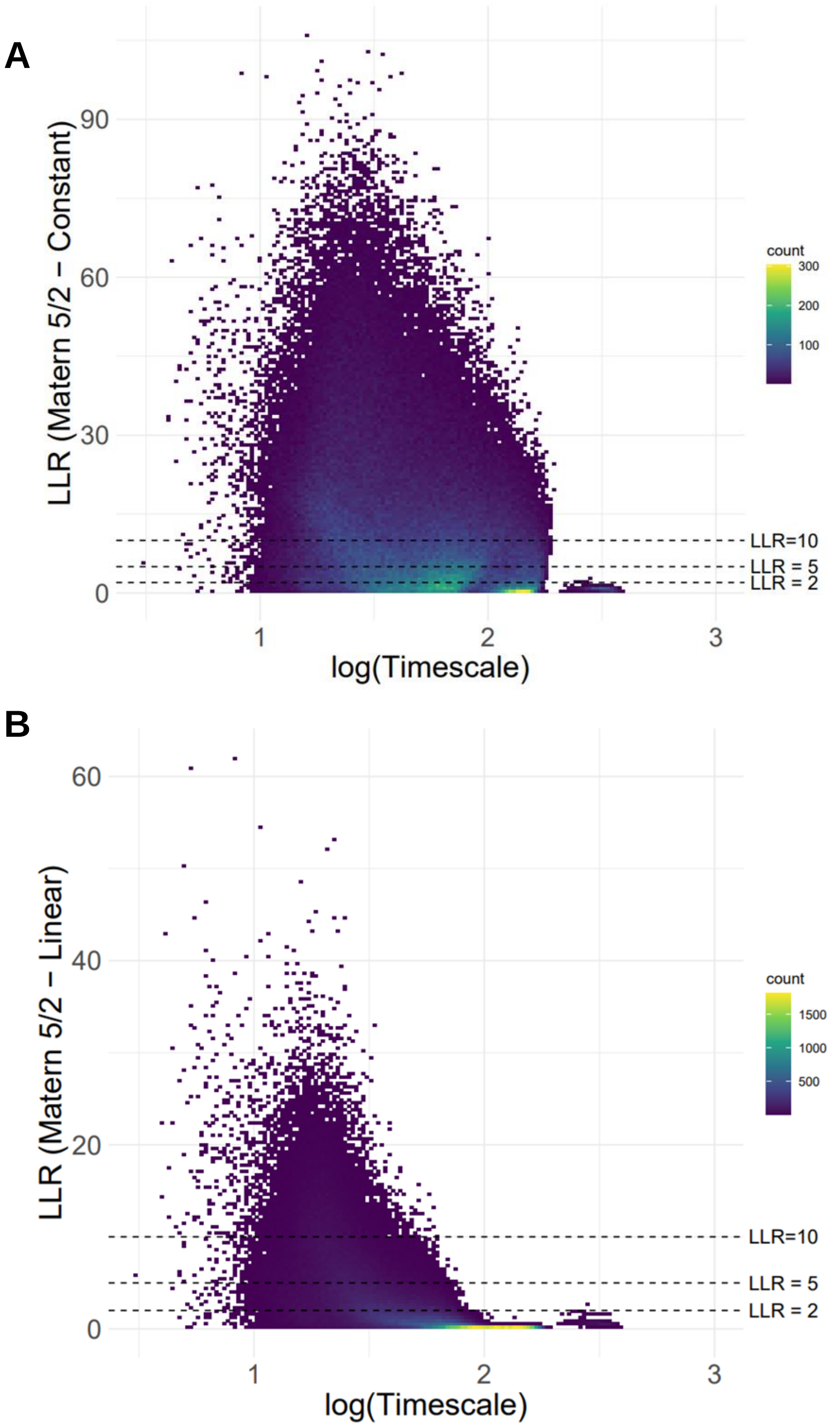

**Supplementary Figure 25 – Nonlinear DNA methylation sites most representative of the module eigengene (“hub sites”).** For each of the six nonlinear modules, shown are the top three DNA methylation sites most highly correlated to the module eigengene.

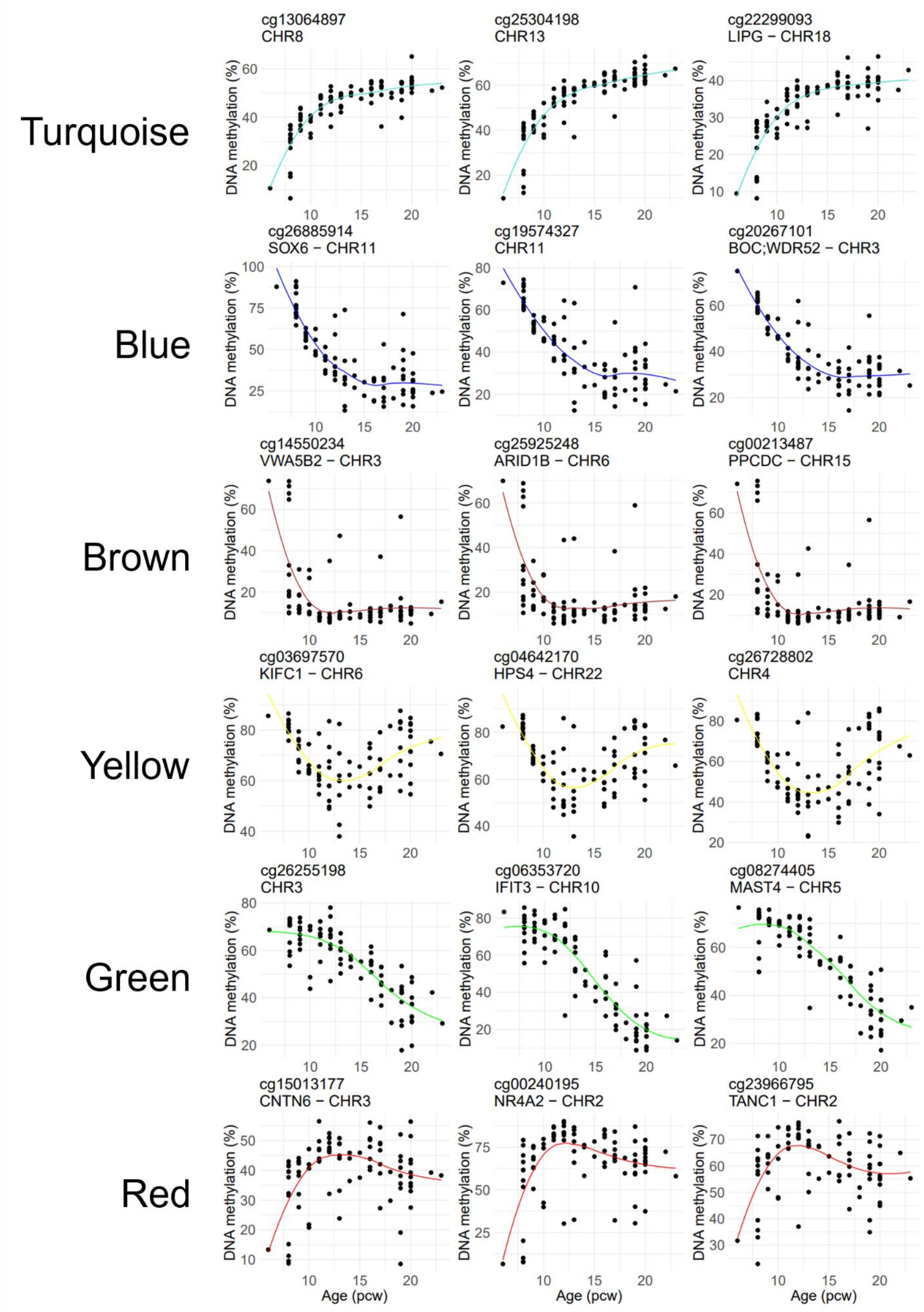
